# The sulfur-related metabolic status of *Aspergillus fumigatus* during infection reveals cytosolic serine hydroxymethyltransferase as a promising antifungal target

**DOI:** 10.1101/2024.03.08.582945

**Authors:** Reem Alharthi, Monica Sueiro-Olivares, Isabelle Storer, Hajer Bin Shuraym, Jennifer Scott, Reem Al-Shidhani, Rachael Fortune-Grant, Elaine Bignell, Lydia Tabernero, Michael Bromley, Can Zhao, Jorge Amich

## Abstract

Sulfur metabolism is an essential aspect of fungal physiology and is known to be crucial for pathogenicity. Fungal sulfur metabolism comprises anabolic and catabolic routes that are not well-conserved in mammals, and therefore can be considered a promising source of prospective novel antifungal targets. To gain insight into the status of the *Aspergillus fumigatus* sulfur-related metabolism during infection we used a NanoString custom nCounter TagSet and compared the expression of 68 key metabolic genes in different murine models of invasive pulmonary aspergillosis, at three different time-points, and a variety of *in vitro* conditions. We identified a set of 15 genes that are consistently expressed at higher levels *in vivo* than *in vitro*, suggesting that they may be particularly relevant for intrapulmonary growth and therefore constitute promising drug targets. Indeed, the role of five of the fifteen genes had previously been empirically validated, supporting the likelihood that the remaining candidates are relevant. In addition, the analysis of the dynamics of gene expression at the early (16h), mid (24h-1) and late (72h) time-points uncovered potential disease initiation and progression factors. We further characterised one of the identified genes, encoding the cytosolic serine hydroxymethyltransferase ShmB, and demonstrated that it is an essential gene of *A. fumigatus* and that it is also required for virulence in a murine model of established pulmonary infection. We further show that the structure of the ligand binding pocket of the fungal enzyme differs significantly from its human counterpart, suggesting that specific inhibitors can be designed. Therefore, *in vivo* transcriptomics is a powerful tool to identify genes crucial for fungal pathogenicity that might encode promising antifungal target candidates.

**AUTHOR SUMMARY:** *Aspergillus fumigatus* is an opportunistic human fungal pathogen that causes devastating chronic and invasive infections in immunocompromised patients. Our arsenal of antifungal drugs to fight this and other fungal pathogens is very limited, partly because of the high similarity between eukaryotic fungal and human cells makes the identification of suitable drug targets a challenging task. Furthermore, targets identified *in vitro* are often not effective *in vivo*, as their action is not relevant for fungal virulence. To address this challenge, we compared the expression profiles of a set of genes involved in sulfur metabolism, a promising source of potential drug targets, in numerous *in vitro* and *in vivo* conditions to identify favourable antifungal candidates. Subsequently, we validated one of the highlighted genes, demonstrating that it is essential for *A. fumigatus* viability and virulence, and that it can likely be targeted by specific inhibitors. Hence, we show the potential of using *in vivo* transcriptomics to identify targets that contribute to virulence, propose various candidates for future studies and present a novel target validated for further antifungal drug development.

## INTRODUCTION

*Aspergillus fumigatus* is a ubiquitous mould that normally lives in the soil feeding on decaying organic matter [1]. During its natural life-cycle *A. fumigatus* produces thousands of spores, also known as conidia, that are disseminated in the air. Because of their high abundance and prevalence, it has been estimated that every human breathes in several hundred of these conidia on a daily basis [2] and, due to their small size (2-3 µm), they have the potential to penetrate deep into the respiratory tract and even reach the lung alveoli [3]. In immunocompetent individuals, this has no consequence as epithelial mucocilliarity and a proper immune response eliminate the spores very efficiently [4]. However, patients with an imbalanced immune response are susceptible to *A. fumigatus*, which can cause a wide range of diseases collectively termed aspergilloses [5]. Chronic and invasive aspergilloses are life-threatening infections with high mortality rates, even in patients under antifungal treatment [6, 7]. First line therapy against aspergilloses is based on the use of azoles, the only class of antifungals that can be orally administered. Worryingly, the number of clinical *A. fumigatus* isolates that are resistant to triazole antifungals is increasing worldwide, which correlates with higher mortality rates [6, 8, 9]. Consequently, new agents to fight this fungal pathogen are urgently required.

Metabolism is central to the virulence of fungal pathogens, and therefore its targeting is considered a promising strategy for the development of novel antifungals [10, 11]. Fungal sulfur metabolism is particularly interesting as it comprises routes and enzymes that are often not conserved in humans, and thus it is considered a propitious source of novel antifungal targets [12]. Indeed, we have previously shown that a proper regulation of sulfur metabolism is crucial for *A. fumigatus* infective capacity [13], demonstrated that biosynthesis of the S-containing amino acids is required for virulence [14], and comprehensively characterised an enzyme in the trans-sulfuration pathway, methionine synthase, as a promising antifungal target [15]. Others have also shown that sulfur is involved in the biosynthesis of essential fungal metabolites [16] and proposed sulfur assimilation and the trans- sulfuration pathway as sources of antifungal targets [17, 18]. Here, we aimed to expand on that knowledge by using *in vivo* transcriptomics to better characterise the fungal sulfur metabolic status during infection of the murine lung, and to use the expression profile of genes to identify those that may be relevant for infection, proposing them as promising candidates for further investigations. However, detection and analysis of the fungal transcriptome during infection is a persistent challenge, as fungal RNA usually represents only between 0.05 and 0.5% of the total RNA isolated from infected organs [19]. To overcome this obstacle, we have used NanoString nCounter, a novel probe-based technology with excellent reproducibility, sensitivity, and specificity when compared to other transcriptomic methods such as microarray hybridisation or RT-qPCR [20, 21]. Its high sensitivity, being capable of detecting mRNA concentrations in the femtomolar range [22], makes this technology ideal for detecting and quantifying fungal RNA during infection. Furthermore, its incomparable specificity is fundamental to prevent any crosstalk with the abundant host RNA. Using this technology, we have assessed the transcription profile of a selection of metabolic genes during infection of the murine lung in two models of immunosuppression and at three time-points post infection, and compared them with the expression profiles under a variety of *in vitro* conditions. By these means we have uncovered interesting aspects of the *in vivo* sulfur metabolic status and identified genes that can be proposed as relevant for growth in the tissues. We have further characterised a promising candidate, the cytosolic serine hydroxymethyltransferase (*shmB*) encoding gene, demonstrating that it is essential for *A. fumigatus* viability as well as for virulence and that it is promising to aim for the development of fungal-specific inhibitors against this target, which can serve to address the emerging global threat of antifungal resistance [23, 24].

## RESULTS

### Linear amplification of Aspergillus fumigatus mRNAs allows gene expression profiling of low input in vivo samples

To determine the transcriptional status of genes related to sulfur metabolism during fungal infection of the murine lung, we developed a custom NanoString nCounter Elements TagSet comprising 68 metabolic genes (51 directly related to S-metabolism and 17 metabolic genes of interest indirectly related with S-metabolism) and 4 housekeeping controls (Table S1). We isolated RNA from *A. fumigatus* propagated under a variety of *in vitro* conditions (Table S2) and from the lungs of infected mice immunosuppressed with two different regimens, the leukopenic and the corticosteroid model (Table S2 and see material and methods). We isolated the lungs of infected mice at three different time-points to investigate gene expression at an early (16h), medium (24h) and late (72h) time point of infection (3 mice per time-point). We further assayed a chronic model of pulmonary aspergillosis *A.* [25] to determine if this model provides a significantly different S-environment (Table S2). A preliminary experiment revealed that fungal RNA could not be reliable detected at 16h and 24h post-infection, which is expected as the fungal burden at those early times is very low. To circumvent this problem, we applied the Nanostring Single Cell Gene Expression protocol (see Material and Methods) to perform linear amplification using Multiplexed Target Enrichment (MTE) primers. To verify that the MTE enrichment does not affect the relative abundance of transcripts, we compared the fold changes between diluted-amplified and non-amplified (undiluted) *in vitro* (-S limiting conditions) and *in vivo* samples (lungs extracted from two leukopenic mice 72h after infection) (Fig 1A). This direct comparison demonstrated that the correlation of the fold changes detected with non-amplified and amplified samples was very high (R^2^=0.95 for mouse 1 and R^2^=0.84 for mouse 2). Therefore, MTE enrichment of low input fungal RNA samples does not affect the relative abundance of transcripts and can be used to investigate gene expression in the mammalian lung at early time-points after infection.

**Figure 1.**
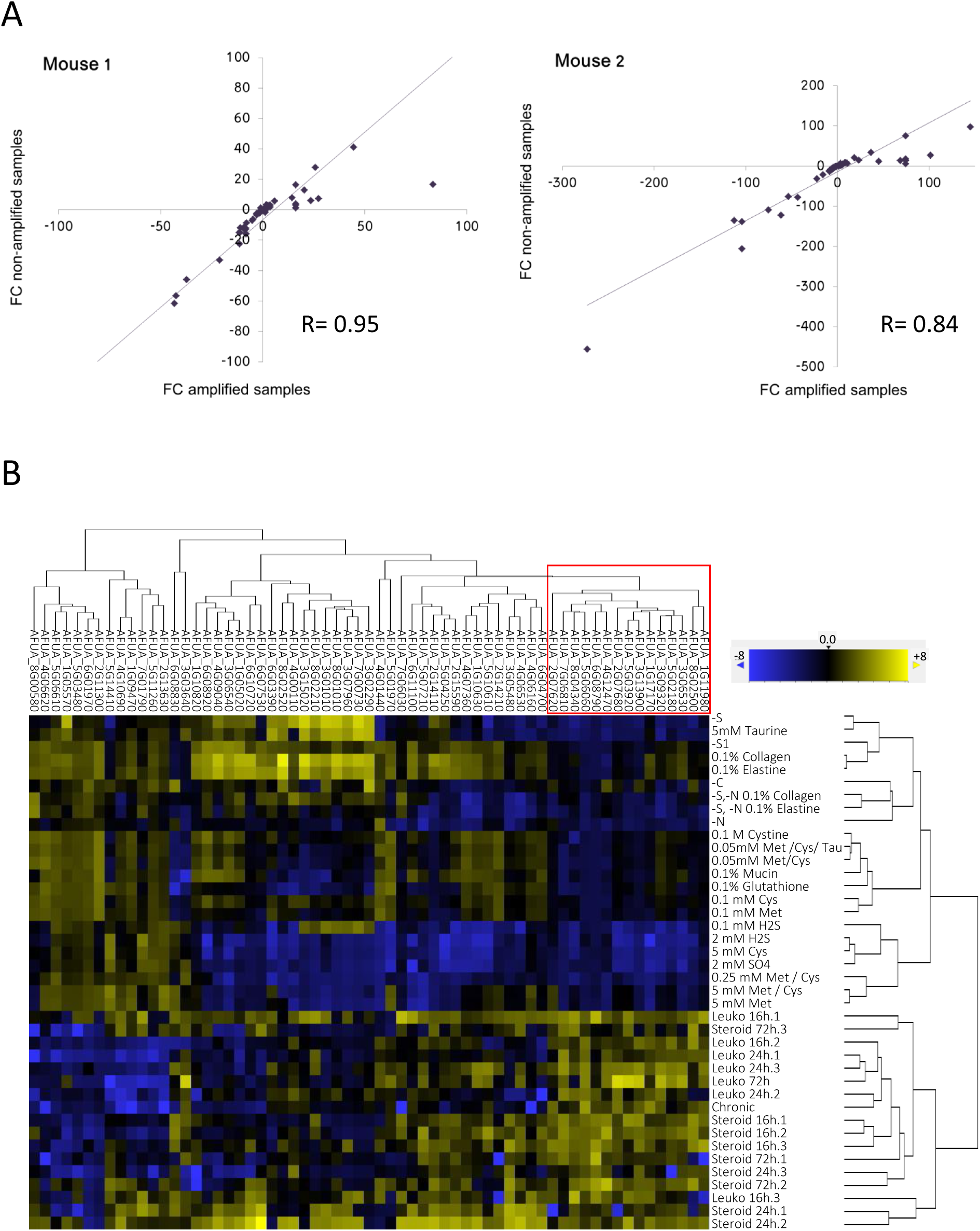
Identification of genes potentially relevant *for Aspergillus fumigatus* virulence using NanoString *in vivo* transcriptomics. A) Validation of the linear amplification performed with the Nanostring Single Cell Gene Expression protocol with Multiplexed Target Enrichment (MTE) primers. The fold changes between diluted- amplified and non-amplified (undiluted) *in vitro* (-S limiting conditions) and *in vivo* samples (lungs extracted from two leukopenic mice 72h after infection) were compared. For both mice, there was a linear relationship between the FCs of non-amplified and amplified samples, demonstrating that amplification did not affect the relative abundance of transcripts. **B)** Hierarchical clustering of all samples formed two distinct branches, one of *in vitro* samples and one with the *in vivo* samples. There were 15 genes (red line) that were always expressed at higher levels in the *in vivo* samples, compared with all the *in vitro* samples.

### The fungal S-related metabolic status in vivo is different from that of in vitro conditions, revealing genes potentially relevant for infection

To analyse the S-related fungal metabolic status *in vivo* and *in vitro*, we hybridized and read our custom NanoString codeset with isolated RNAs or amplified cDNAs (Table S2). The obtained raw data was subjected to quality control, which revealed that the samples obtained from two leukopenic mice 72h had not run properly: they were borderline for the recommended threshold of overall number of counts (10,000) and did not reach the minimum values for the positive counts reads (Fig. S1A and S1B). Consequently, these samples were excluded from all analyses. In addition, inspection of the normalized data (see Material and Methods for details) revealed that five genes (AFUA_3G06492, AFUA_1G06940, AFUA_5G08600, AFUA_4G03950 and AFUA_6G00760) had a very low number of counts (<20) in all samples. This could be due to the genes not being expressed at all or at very low levels in the tested conditions, or to failure of the NanoString designed probes to detect these mRNAs. Accordingly, these genes were also excluded from all downstream analyses.

We first performed an unsupervised hierarchical clustering of all samples analysed (Fig. 1B). By this stratification, the *in vivo* samples clearly clustered separately from the *in vitro* samples, indicating that the fungal S-related metabolic status during infection is significantly different from that during growth in diverse experimental conditions This suggests that the lung provides a unique environment in terms of sulfur metabolism that is clearly distinct from defined culture media, despite our efforts to provide potentially relevant sulfur sources. Noteworthy, this also included the two S-depleted samples, which indicates that the mammalian lung is not sulfur limiting. Based on this, we hypothesised that genes which are consistently more highly expressed *in vivo*, compared to in the great variety of tested *in vitro* conditions, may be particularly relevant for the fungal capacity to grow in the lung tissues, and therefore for fungal virulence. We detected a cluster of 15 genes which were always expressed more *in vivo* than *in vitro* (Fig. 1B, red box, Table 1). Interestingly, we and others have already demonstrated the importance of at least five of those genes for *A. fumigatus* virulence, and a further six genes had already been proposed to be potentially relevant (Table 1 and see discussion), supporting the rationale that the uncharacterized genes might be valid candidates for further analyses.

**Table 1.** Genes that consistently showed higher expression *in vivo* compared to any of the tested *in vitro* conditions.

| GENE ID | DESCRIPTION | VIRULENCE |
| --- | --- | --- |
| AFUA_1G11980 | Glutamine amidotransferase (HisfF/H) | Suspected [26] |
| AFUA_8G02500 | Glutathione S-transferase | Suspected [27] |
| AFUA_3G06530 | ATP sulfurylase (sC) | Unknown |
| AFUA_5G02180 | Cysteine synthase (CysB) | Proven [14] |
| AFUA_3G09320 | Serine hydroxymethyltransferase (SmhB) | This work |
| AFUA_1G17170 | Taurine dioxygenase | Unknown |
| AFUA_3G13900 | Glutamate-cysteine ligase (Gcs1) | Suspected [27] |
| AFUA_5G03920 | bZIP transcription factor (HapX) | Proven [28] |
| AFUA_2G07680 | L-ornithine N5-oxygenase (SidA) | Proven [29] |
| AFUA_4G12470 | bZIP transcription factor (CpcA) | Proven [30] |
| AFUA_6G08790 | C6 transcription factor (PrnA) | Unknown |
| AFUA_5G06060 | SCF-complex subunit (SkpA) | Suspected [31, 32] |
| AFUA_8G04340 | cystathionine gamma-lyase (MecB) | Proven [33] |
| AFUA_7G06810 | L-amino acid oxidase (LaoA) | Suspected [34] |
| AFUA_2G07620 | Cystathionine beta-synthase (mecA) | Suspected [33] |

### Fungal sulfur related in vivo metabolism is influenced by the model of immunosuppression and the course of infection

To investigate if the fungal S-related metabolic status is influenced by the model of immunosuppression, we performed hierarchical clustering of all the *in vivo* samples (Fig. 2A). This clustering showed that the fungal S-related status is quite variable *in vivo* (notice that the dynamic range of expression are much smaller than when comparing *in vitro* and *in vivo* samples in Fig. 1B), and is partially dependent on the model of immunosuppression, as 5 out of the 8 leukopenic mice aggregated in a separated cluster. Interestingly, the chronic model clustered in this separated branch, suggesting that the fungal S-related metabolic status in this type of infection is more similar to the status in the leukopenic model than in the steroid-induced one. Nevertheless, some samples from leukopenic mice interleaved with the samples from steroid-treated mice, demonstrating that the model of immunosuppression is not the only factor that determines the fungal metabolic status.

**Figure 2.**
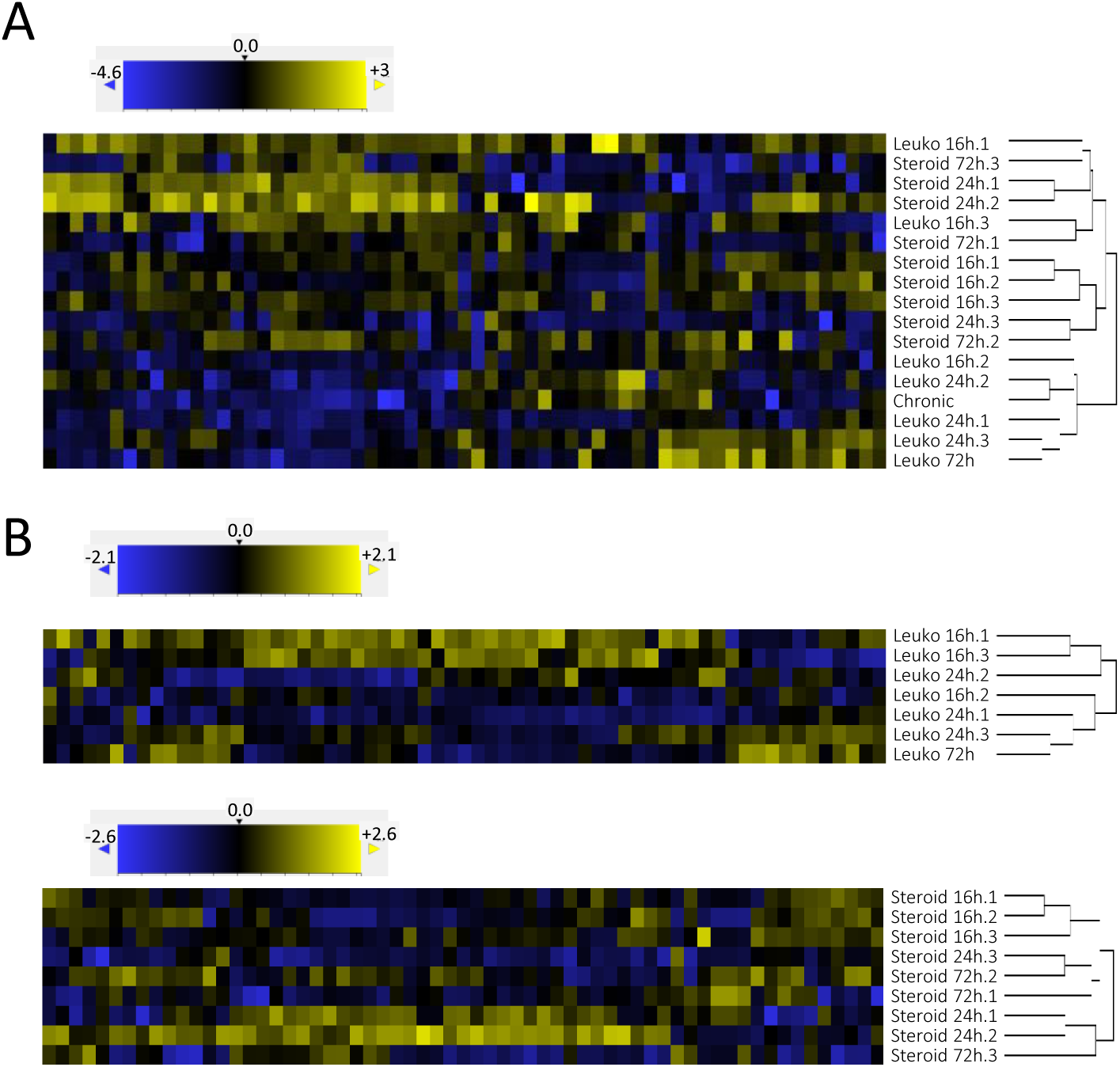
The fungal S-related sulfur metabolism *in vivo* is influenced by the model of suppression and the time of infection. **A)** Hierarchical clustering of all *in vivo* samples created two major branches, one almost exclusive for leukopenic samples, which only included the chronic model sample. The second major branch was divided again in two branches, one exclusive for steroid samples and the other mixed with leukopenic and steroid samples. Therefore, the model of suppression influences but is not the only factor that determines the fungal metabolic status. **B)** Separated hierarchical clustering of leukopenic or steroid samples partially aggregated by the time of infection. Therefore, the fungal S-related metabolism is influenced *in vivo* by the status of infection.

To investigate to what extent the fungal S-related metabolic status is influenced by the course of infection, we performed hierarchical clustering of the samples from leukopenic and steroid-treated mice separately (Fig 2B). In both models of suppression there was a partial aggregation of samples according to the time point post infection, suggesting that the course of infection is mirrored by the S-related metabolic status of the fungal pathogen.

In conclusion, the model of immunosuppression as well as the time of duration of infection appear to influence the status of the fungal S-related metabolism. However, these variables alone or combined cannot explain the variances among samples, indicating that other factors must be involved. For instance, it is to be expected that the exact anatomic location of the foci of fungal growth would account for different microenvironments that in turn affect the fungal S-related metabolism.

### Temporal dynamics of gene expression in vivo versus in vitro reveals the existence of potential disease initiation and progression factors

Since we found that the S-related metabolic status correlates with the time period post infection, we reasoned that some genes may be expressed at different levels as the fungal infection progresses. To investigate this hypothesis, we first examined the fold change of the (geometric) mean expression of all genes in each time-point with respect to the corresponding 16h condition (Fig. 3A). This analysis revealed that indeed many genes change the level of expression during the course of infection: transcription of several genes seemed to consistently increase, a few decreased expression as infection progressed, and some appeared to increase or decrease expression at mid-infection (24h) before returning to initial levels in late infection (72h). To explore whether any of these genes that show temporal dynamic expression could be particularly important for *in vivo* growth, we analysed the fold change of all genes in each *in vitro* condition versus each of the leukopenic (Fig. S2A) or steroid-treated (Fig. S2B) samples grouped by the time after infection. As expected, we identified some genes which showed a temporal variance *in vivo* compared with *in vitro*. In detail, in leukopenic mice we observed that genes *metAT* (AFUA_2G13630), *cdoB* (AFUA_5G14410), *cysAT* (AFUA_1G09470), *isa1* (AFUA_4G10690) and *hisB* (AFUA_6G04700) had higher transcript levels *in vivo* at early (16h) than at mid (24h) or late (72h) time points after infection (Fig. 3B and Fig. S2A orange boxes), suggesting that these genes may be particularly important for the initiation of infection, but possibly not as relevant for progression and maintenance of infection. In contrast, *cdoA* (AFUA_1G05570) had higher *in vivo* transcription during advanced infection (Fig. 3C and Fig. S2A purple box), suggesting that this gene may be relevant for growth in progressively invaded tissue. The genes, *sD* (AFUA_1G10820), *metR* (AFUA_4G06530) and *hapX* (AFUA_5G03920) displayed higher expression at early (16h) and late (72h) time after infection (Fig. 3D and Fig. S2A black box), suggesting that they may be particularly important to initiate infection and during established infection, but that they are not crucial for infection to progress.

**Figure 3.**
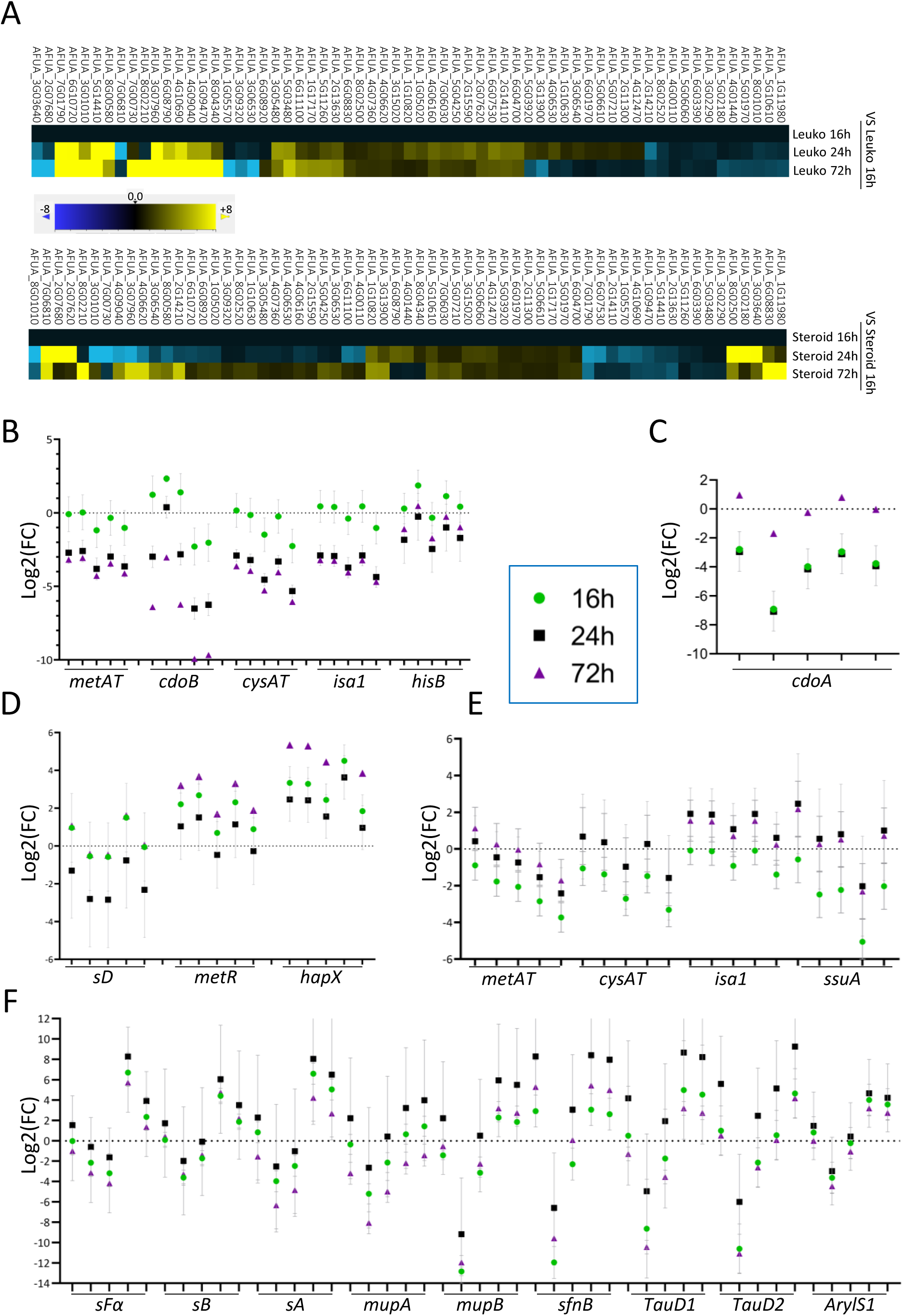
Temporal dynamics of gene expression reveals the existence of potential initiation and disease progression factors. **A)** Fold change of expression of genes in each time-point with respect to the correspondent 16h condition revealed that most genes changed expression during the course of infection. The geometric mean value for each time-point was used. **B-F** panels display the Log2 fold change (FC) of genes that showed temporal dynamic expression in five randomly selected representative *in vitro* condition (in order: -N, -S, 0.1 mM Cys, 2 mM SH2 and 5 mM Met) versus *in vivo* conditions. All graphs display the mean and the standard error of the mean (SEM). **B)** These five genes always showed higher expression at early (16h) than at mid- or late (24h or 72h) infection of leukopenic mice when compared with the *in vitro* conditions. **C)** In leukopenic mice, the gene *cdoA* consistently showed the highest level of expression at late infection, when compared with the *in vitro* conditions. **D)** These three genes always showed higher expression at early and late (16 and 72h) than at mid- (24h) infection of leukopenic mice, when compared with the in vitro conditions. **E)** The four genes consistently showed higher expression at mid- and late infection than at early infection, when compared with the *in vitro* conditions. **F)** These 9 genes always showed the highest level of expression at mid- (24h) than at early or late infection, when compared with the *in vitro* conditions.

Similarly, in steroid treated mice, we observed that genes *metAT* (AFUA_2G13630), *cysAT* (AFUA_1G09470), *isa1* (AFUA_4G10690) and *ssuA* (AFUA_7G01790) had higher expression at mid- (24h) and late (72h) times after infection (Fig. 3E and Fig. S2B purple box), indicating that they may be more relevant for progression and maintenance of infection than for its initiation. Finally, genes *sFα* (AFUA_6G08920), *sB* (AFUA_1G05020), *sA* (AFUA_3G06540), *mupA* (AFUA_4G09040), *mupB* (AFUA_7G00730), *sfnB* (AFUA_8G01010), two taurine dioxygenase encoding genes (AFUA_3G0796 and AFUA_3G01010) and one arylsulfatase encoding gene (AFUA_8G02520) had higher expression *in vivo* at mid-infection time (24h) (Fig. 3G and Fig. S2B orange boxes), suggesting that these genes may play a role in this model for the progression of infection.

We therefore propose that the temporal dynamics of *in vivo* transcript levels may be useful to identify and differentiate fungal factors of disease initiation and progression, defined as products that are essential for the initiation of infection or that facilitate persistence and continued disease progression [35]. Although this hypothesis needs to be validated further by confirming the role of some of our identified candidates as true disease initiation or progression factors in appropriate models of infection, we emphasise the potential of *in vivo* transcriptomics at distinct stages of infection to detect such initiation and progression factors, as has already been proposed [35].

### Validation of an identified gene, encoding a serine hydroxymethyltransferase (SHMT), as a promising target for antifungal drug development

To demonstrate the value of the NanoString technology in identifying genes relevant for *A. fumigatus* virulence, and aiming to validate a novel drug target, we decided to further investigate the serine hydroxymethyltransferase encoding gene (*shmB,* AFUA_3G09320), which was consistently expressed at higher levels *in vivo* compared with *in vitro* conditions (Fig. 1B). *A. fumigatus* genome encodes two paralogues of serine hydroxymethyltransferase, ShmA (annotated as AFUA_2G07810, not included in the NanoString analysis) and ShmB. The ShmB protein contains 471 amino acids (aa) and ShmA is 537 aa in length, and both share an identity of 61.9% and similarity of 79.9% (Fig. S3A). Interestingly, ShmA has a long N-terminus region that is absent in ShmB (Fig. S3A), which prompted us to investigate the subcellular localisation of the proteins using three different public softwares: DeepMito [36], PProwler [37], and TargetP [38]. In contrast to the current annotation in FungiDB and NCBI databases, all three algorithms predicted a mitochondrial localisation for ShmA and a cytosolic for ShmB. Indeed, when we aligned the *A. fumigatus* proteins with their *Saccharomyces cerevisiae* orthologues, we observed that the yeast mitochondrial enzyme SHM1 also had a longer N-terminus in comparison to the cytosolic proteins (Fig. S3B).

Therefore, aiming to clarify the subcellular localisation of ShmA and ShmB in *A. fumigatus* we tagged the proteins at their C-terminus with the fluorescent YFP-derivative Citrine to express them under the control of their native promoters. In line with our predictions, we found that the ShmB-Citrine protein localises throughout the cytoplasm, with a similar pattern to a control strain that expresses Citrine under the control of the strong *gdpA* promoter (Fig. 4A). Regrettably, ShmA-Citrine could not be detected, possibly reflecting very low or no expression of the encoding gene under steady-state conditions, which would be supported by its dispensability under standard laboratory conditions (see below).

**Figure 4.**
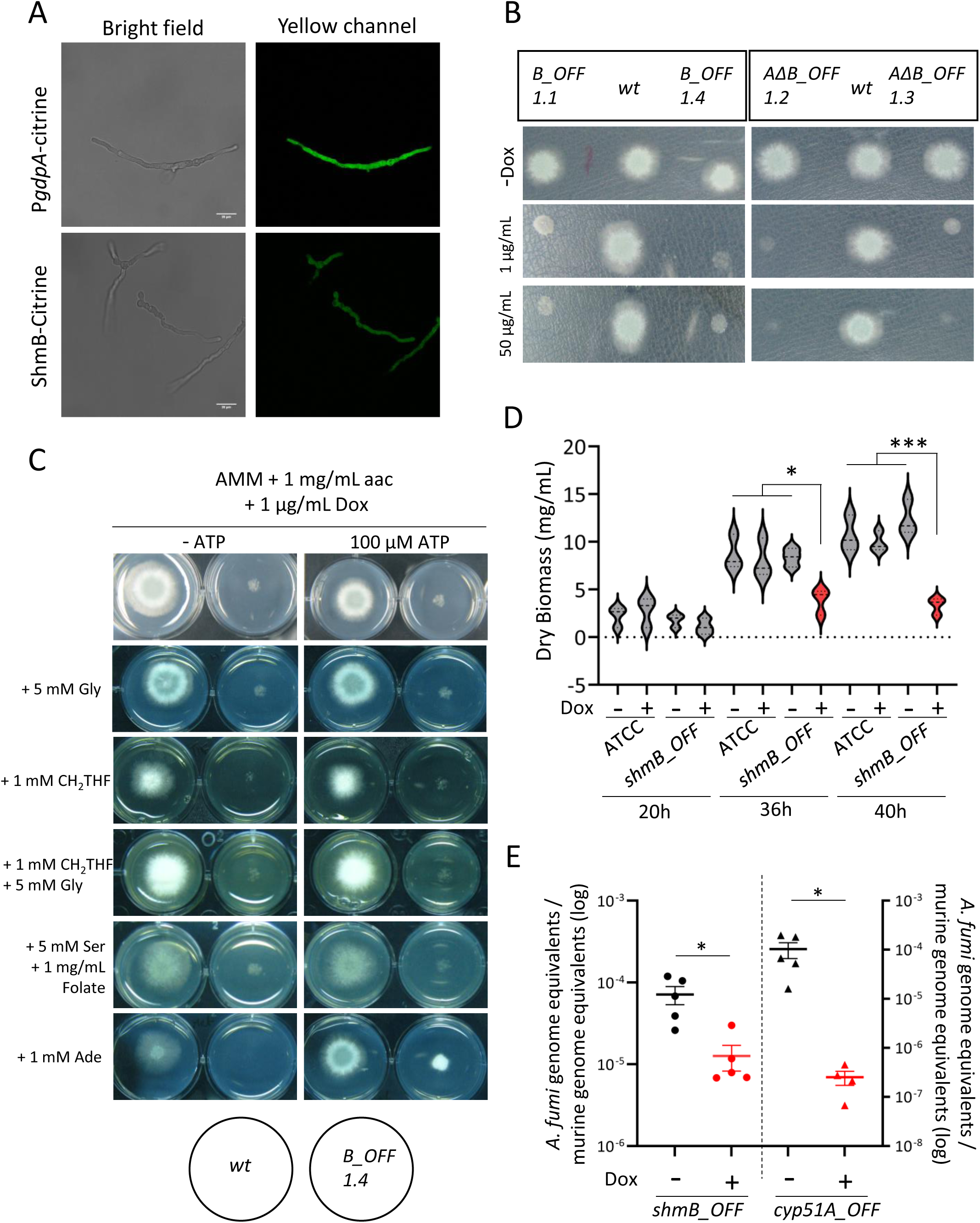
The cytosolic serine hydroxymethyltransferase is essential for viability and virulence in *Aspergillus fumigatus*. **A)** A ShmB-Citrine tagged proteins localised diffused throughout the cytoplasm, similarly as the control P*gdpA*-Citrine strain.**B)** A Phenotypic assay with two independent clones of the *shmB_tetOFF* and *ΔshmA shmB_tetOFF* mutant strains. In the presence of doxycycline (Dox) the mutants did not grow, demonstrating that *shmB* is essential in *A. fumigatus*, and that ShmA cannot cover its function. Plates were incubated for 48h at 37 °C. The experiment was repeated twice independently. **C)** Attempting to reconstitute *shmB_tetOFF* growth in restrictive conditions (+Dox), media was supplemented with various metabolites which could potentially be limited in the absence of serine hydroxymethyltransferase activity. The only condition that could partly reconstitute growth, not at wild-type level, was supplementation with ATP and adenine (Ade). Plates were incubated for 48 h at 37 °C. The experiment was repeated twice independently. **D)** Dox (10 µg/mL) was added or not to 12h grown submerged mycelia of the *shmB_tetOFF* and its parental strains. Measurements of fungal biomass at 20h, 36h and 40h post-inoculation showed a significant reduction of growth after downregulation of *shmB.* The experiment was repeated three times independently, with three technical replicates. The results were analysed using a one-way ANOVA with Tukey’s multiple comparisons test. * = *p*<0.05 and ***=*p*<0.001 **E)** Fungal burden in the lungs of leukopenic mice treated or untreated with Dox. Downregulation of *cyp51A* (encoding the target of the azoles) expression caused a significant reduction in burden (*p*=0.0159 in a nonparametric Mann Whitney test and *p*=0.0476 in an unpaired *t*-test with Welch’s correction). Similarly, downregulation of *shmB* triggered a significant decrease in fungal burden (*p*=0.0159 in a nonparametric Mann Whitney test and *p*=0.0288 in an unpaired *t*-test with Welch’s correction). Therefore, ShmB appears to be a valid antifungal target. Each point represents an independent mouse (biological replicate) and the graph shows the mean and the standard of the mean (SEM). Comparisons were made using a Mann-Whitney non-parametric *t*-test.

To investigate the potential relevance of ShmA and ShmB for fungal virulence, we initially attempted to construct single null mutants deleted for each encoding gene. We successfully constructed a *ΔshmA* mutant, which showed no phenotype on regular laboratory conditions, growing perfectly on rich Sabouraud and defined AMM media (Fig. S3C). In contrast, we were unable to generate a *ΔshmB* mutant, suggesting that this might be an essential gene in *A. fumigatus*. We therefore constructed two mutant strains in which we placed the *shmB* coding region under the control of the *tetOFF* module [39, 40], one in the wild-type background and one in the *ΔshmA* background. The *tetOFF* system allows downregulation of transcription of the gene of interest, in this case *shmB*, upon supplementation of doxycycline (Dox) [39]. Addition of Dox inhibited growth of both strains (*shmB_tetOFF* and *ΔshmA- shmB_tetOFF*) on AMM solid plates (Fig. 4B), proving that *shmB* is essential for viability independently of its paralogue *shmA*. Gene essentiality is a conditional phenomenon, whereby particular growth/environmental conditions may overcome the disturbances derived from the gene deficiency [40–43]. Therefore, we attempted to reconstitute growth of the *shmB_tetOFF* strain in the presence of Dox by supplementing compounds that could potentially be depleted in the absence of ShmB (Fig. 4C). We have previously observed that disturbances in the folate cycle causes a need of amino acid supplementation [44], therefore, we assayed all phenotypic conditions on AMM without nitrogen (AMM-N, to diversify and increase the presence of permeases in the membrane that ensure uptake of amino acids and other compounds) supplemented with 1 mg/mL of all proteinogenic amino acids. Additionally, we tried to reconstitute growth by supplementing the media with 5,10- methylenetetrahydrofolate (CH2-THF) and glycine (separately and combined), or with folic acid and serine (substrates/products of the bidirectional enzymatic reaction), or with adenine (as blockade of the folate cycle might impair purine biosynthesis [45]). In addition, we tested adding ATP to all those supplements, as we previously showed that impairing the folate cycle *via* blocking the activity of methionine synthase impacts cellular energetics in *A. fumigatus* [40]. The only condition that could partly reconstitute growth, albeit not to wild-type levels, was supplementation with adenine and ATP (Fig. 4C). None of the supplement combinations reconstituted growth of the *shmB_tetOFF* strain under restrictive conditions (Fig. 4C). Therefore, either *shmB* is a truly essential gene, or the conditions to overcome its essentiality are complex and yet unknown. Yet, based on these results *shmB* constitutes a promising drug target candidate, and consequently we decided to test its value in an *in vivo* infection model.

We have recently proposed that potential drug targets should be validated in models of established infection, which is particularly relevant for essential genes (to confirm that essentiality cannot be overcome during active growth in the host tissues) [40]. To this aim, we initially confirmed that Dox addition very strongly turns down *shmB* expression in growing mycelium (Fig. S3D) and that this downregulation causes a significant decrease in the growth capacity of 12h grown submerged mycelia, as measured by fungal biomass (Fig. 4D). Therefore, we assayed our optimised protocol to investigate targets in established *A. fumigatus* invasive pulmonary infections with the *tetOFF* system [40]. We infected leukopenic mice with 2×10^5^ conidia of the *shmB_tetOFF* strain, started Dox treatment 16h after infection and measured fungal burden in the lungs 72h after infection. As control we also infected mice with the previously validated strain *Δcyp51B-cyp51A-tetOFF* [40], which expresses *cyp51A*, the gene that encodes the target of the azole antifungals, under the control of the same *tetOFF* system. Therefore, this control strain serves to corroborate that downregulating a validated target (azoles are the gold standard therapy for invasive pulmonary aspergillosis) causes a significant reduction in fungal burden, as indeed we detected (Fig. 4E). Interestingly, we found that downregulation of *shmB* in the established infection model triggered a significant reduction in fungal burden (Fig. 4E). Hence, ShmB appears to be a valid target for the treatment of aspergillosis infections.

### Structural-based analysis of ShmB druggability

The human serine hydroxymethyltransferase (SHMT) is considered a promising target for anti-cancer drug development [46, 47] and accordingly various inhibitors have been already developed (for instance, see references in [48]). Therefore, we decided to test the antifungal potential of one of such inhibitors of the human SHMT, Hit-1 [49], on *A. fumigatus*. We performed broth microdilution assays incubating the fungus in RPMI-1640 medium in the presence of increasing concentrations of Hit-1 (from 2 µg/mL=4.29 µM to 512 µg/mL=1.1 mM), and did not detect any antifungal activity of this compound (Table 2). As this could be due to the incapacity of Hit-1 to penetrate fungal cells, we also tested Hit-1 antifungal activity in the presence of colistin. This antibiotic has been shown to permeabilise *Candida albicans* cells [50], and here we have confirmed that it also permeabilises *A. fumigatus* swollen conidia to the membrane impermeable dyes FITC (fluorescein isothiocyanate) and Zombie-Aqua (Fig. S4). However, Hit-1 did also not inhibit *A. fumigatus* growth in the presence of colistin (Table 2).

**Table 2.** Broth microdilution assays of colistin and Hit-1 showed that neither of these compounds inhibit growth at the tested concentrations.

|  | Minimum Inhibitory Concentration (MIC) |
| --- | --- |
| **Colistin** | > 64 µg/mL |
| **Hit-1** | > 512 µg/mL |
| **Hit-1 + 2 µg/mL Colistin** | > 512 µg/mL |
| **Hit-1 + 4 µg/mL Colistin** | > 512 µg/mL |

As our previous results with the *tetOFF* promoter demonstrated that ShmB is essential for *A. fumigatus* viability, we suspected that the inhibitor Hit-1, designed against the human enzyme, may not be effective against the fungal enzyme. This led us to hypothesise that it should also be possible to design fungal specific inhibitors, not active against the human enzyme. To substantiate this hypothesis our initial step involved identifying the main druggable binding pocket of the human SHMT based on its crystal structure as retrieved from the Protein Data Bank (accession number PDB ID 1bj4), using a target-based *in silico* approach. We conducted two independent analyses employing the DrugRep [51] and the PockDrug [52] engines. The PockDrug analysis yielded ten possible binding pockets, with only six of them possessing a Druggability Probability score exceeding 0.5 (Fig. 5A). Cross-referencing with the results from DrugRep revealed a single common pocket from both predictions: pocket A (Fig. 5A) (Gln44 Ala383 Ile51 Glu374 Phe56 Ser381 Ala379 Glu69 Val46 Arg43 Cys384 Ala407 Gly47 Glu49 Ile382 Arg465 Glu378 Glu54 Asn55 Leu48). Next, we investigated whether this predicted binding pocket is preserved in the *A. fumigatus* orthologue. To this purpose, we constructed a homological model of the fungal protein using the *in silico* protein modelling AI AlphaFold [53]. Although the similarity between the sequences is reasonably high (74.9%), indicating the protein is well preserved crossed the species, there are some mutations in the amino acids within the key areas associated to the hypothetical binding pocket (highlighted in red in Figure 5B). A close comparison revealed that such mutations likely altered the conformation of the neighbouring amino acid residues hence resulting in a different topology of pocket A compared to human SHMT, making it much shallower and smaller (Figure 5C). In particular, the alteration in the position of Arg263 (Arg255 in *A. fumigatus* SHMT) has formed a barrier resulting in the fungal pocket being almost half the size of the human pocket A.

**Figure 5.**
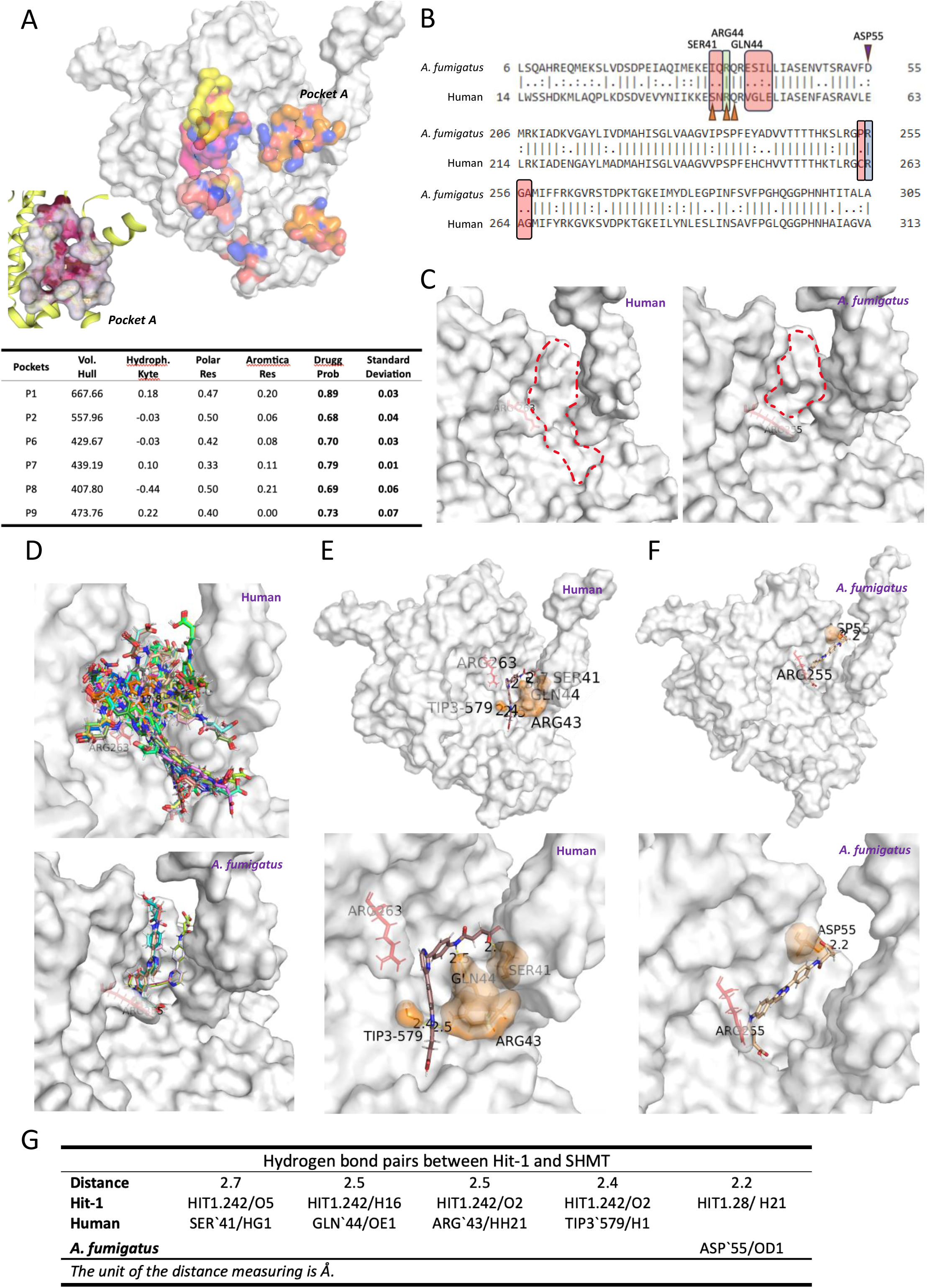
An inhibitor of the human SHMT does not bind properly to the fungal enzyme’s ligand binding pocket. **A)** Binding pockets with draggability score above 0.5 predicated by DrupRep highlighted in human SHMT1 1bj4, with ‘Pocket A’ Zoomed in (left corner). **B)** Alignment of the amino acid sequences (partial) of the *A. fumigatus* (ShmB) and human (SHMT1) cytosolic SHMT enzymes. Different amino acids (highlighted in red rectangles) were detected surrounding the key residues Arg263 in SHMT1 (Arg255 in ShmB, highlighted in blue rectangle); and Arg43 (Arg35 in ShmB, highlighted in green rectangle). Orange arrows indicate SHMT1 residues forming hydrogen bonds with Hit-1 and purple arrow indicates ShmB residue forming hydrogen bond with Hit-1. **C)** Predicted binding pockets comparison between SHMT1 and ShmB with Arg263 (Arg255) highlighted in red. **D)** Hit-1 binding motifs suggested by SwissDock. **E)** Detailed interactions formed between Hit-1 and SHMT1. **F)** Detailed interactions formed between Hit-1 and ShmB. **G)** List of hydrogen bond pairs between Hit-1 and SHMT1 and ShmB.

Subsequently, we performed *in silico* docking using the SwissDock engine [54, 55] employing the Hit- 1 inhibitor (IC50 = 0.53 µM) [49] against pocket A of the human protein. This resulted in several promising binding motifs occupying the entire pocket, as visually represented in Figure 5D. For instance, in one of the top binding positions, HIT1.242 (ΔG = -7.57 kcal/mol; FullFitness = -2433.57 kcal/mol), Hit-1 nestled itself deep into the pocket, demonstrating hydrophobic interactions with the pocket (Fig. 5E). Additionally, it also formed four hydrogen bonds distributed through the molecule backbone (Ser‘41, Gln‘44, Arg‘43 and Tip3‘579; Fig. 5G), anchoring the ligand neatly into the pocket (Fig. 5E). Subsequent *in silico* docking between Hit-1 and the fungal orthologue pocket A suggested that Hit-1 could not bind as effectively. Only two binding motifs were suggested, both restricted to the smaller *A. fumigatus* pocket (Figure 5D). Upon a closer inspection of the top binding motif, HIT1.28 (ΔG = -6.87 kcal/mol; FullFitness = -2143.87 kcal/mol; Fig. 5G), it is clear that the ligand is not embedding deep into the pocket, indicating less hydrophobic interactions with the target. There is only one hydrogen bond suggested between Hit-1 and the target (ASP‘55; Fig. 5F), indicating a less favourable binding compared to the human version of the protein. It also appeared that the altered Arg255 indeed acted as a barrier, blocking Hit-1 from extending to the region proximal to Arg43 (Arg35 in *A. fumigatus* SHMT) and forming any additional hydrogen bonds (Fig. 5F). Therefore, it appears that Hit-1 is specific to human SHMT as it cannot bind properly to the fungal orthologue, which would agree with the observed absence of growth inhibition on *A. fumigatus* (Table 2). Similarly, we believe that inhibitors that bind to the fungal, but not the human, enzyme pocket can be designed, and thus we propose that the development of novel fungal-specific inhibitor agents against the promising SmhB antifungal target is feasible.

## DISCUSION

New antifungal agents are urgently needed, as resistance to the current drugs is widespread [56–58], posing a great threat to human health [24]. In the endeavour of developing novel and efficient chemotherapy, the selection of valid drug targets is crucial [59]. In the last few years, a variety of innovative approaches have been applied to identify suitable target candidates. For instance, comparative genomics [60], computer modelling [61], *in silico* analysis of protein interactions [62], screening of deletion libraries [63], including competitive fitness assays [64], screening of regulatable libraries [43, 65], construction of transposon insertion libraries [66, 67], or machine learning approaches [68] have been employed to identify antifungal drug targets. In this regard, *in vivo* transcriptomics has also been used frequently to understand fungal pathogenicity [34, 69–72], albeit its potential to identify suitable antifungal targets still encounters some scepticism. In a seminal study, McDonagh and colleagues [34] compared the transcriptome obtained from conidia germinating in murine lungs with germlings obtained from *in vitro* conditions expected to match the *in vivo* environment. This work could confirm that *A. fumigatus* encounters nutrient (particularly iron and nitrogen) limitation as well as alkaline and oxidative stress during initiation of infection, providing foundational cues about the conditions encountered by the fungal pathogen in the mammalian lung. However, this study needed to use pools of 24 mice per sample, which limited the number of conditions that could be tested (for instance, only one model of immunosuppression was assayed), could only analysed the early initiation of infection (12-14 hours) and had to recover the fungal material from bronchoalveolar lavages. These limitations were mainly due to the challenge of recovering sufficient fungal RNA material from the infected tissue for the subsequent analysis. In a more recent study, NanoString analysis was employed to identify *A. fumigatus* transcriptional factors which are highly expressed or strongly up-regulated during growth in the murine lung tissues, and experimentally proved that two of them are relevant in the context of infection [73]. Here we reasoned that by comparing the transcription levels of genes related to sulfur metabolism in the tissues versus a variety of *in vitro* S-sources could reveal gene products that are specifically upregulated *in vivo*, and may therefore be relevant for the infection process and constitute promising antifungal targets.

In our analysis of genes related with sulfur metabolism, we have identified 15 genes that are consistently expressed at higher levels in the murine lungs than in all tested *in vitro* conditions. At least 5 of those genes have been previously proven to be important for *A. fumigatus* virulence, and additionally 6 genes have been suspected to be relevant as well (Table 1). We previously showed that AFUA_5G02180, encoding cysteine synthase (CysB), is important for *A. fumigatus* virulence, as a cysteine auxotroph had reduced virulence in a murine model of infection [14]. We have also demonstrated that AFUA_8G04340 encodes cystathionine-γ-lyase (MecB), the major persulfidating enzyme in *A. fumigatus*, which is relevant for fungal virulence [33]. The genes AFUA_2G07680, encoding L-ornithine-*N* ^5^-monooxygenase (SidA), which catalyses the first committed step of siderophore biosynthesis, and AFUA_5G03920, encoding the master transcriptional regulator for adaptation to iron limitation (HapX), have been shown to be crucial for virulence [28, 29]. Finally, the gene AFUA_8G04340, encoding the transcriptional activator of the cross-pathway control system of amino acid biosynthesis (CpcA), was also reported to contribute to *A. fumigatus* virulence [30]. Therefore, we propose that the remaining genes are also likely to be important for *A. fumigatus* virulence and thus constitute good candidates to validate experimentally. It is noteworthy that genes that we have characterized and shown to be dispensable for virulence (AFUA_1G05020 -*sB*- and AFUA_2G15590 -*sF*- [14]) were not detected in this analysis, further supporting the hypothesis that the genes identified may indeed be relevant. Nevertheless, it also needs to be mentioned that several genes that have been described as relevant for virulence in other studies were not identified as universally important in our analysis (for instance, AFUA_6G04700 -HisB- , AFUA_4G06530 -MetR- or AFUA_2G14210 - Ilv3A/IlvC- [13, 26, 74]), implying that our approach is not able to detect all genes relevant for infection. Yet, our analysis has identified some of these genes as potential disease initiation or progression factors, which opens the question of whether their relevance for infection might be time-dependent.

In fact, our temporal analysis has revealed various potential disease initiation and progression factors. These are defined as products that are required for the initiation of infection or for the persistence and progression of disease [35]. Differentiating these actions is paramount for drug discovery, as it would not be adequate to develop a drug to treat established infections that targets an initiation factor, and similarly, it would not be appropriate using a drug that targets a progressing factor as prophylaxis. Therefore, discerning disease initiation and progression factors would be important to better understand the infective capacity of *A. fumigatus* and to expand our reservoir of potential antifungal targets. In fact, progression factors, which are usually not identified in classic virulence studies, are regarded as valuable targets, as antimicrobials are often employed to treat established infection [35]. In this respect, we previously negated a role of *sFα* (AFUA_6G08920) and *sB* (AFUA_1G05020) in *A. fumigatus* virulence by infecting leukopenic mice with null mutants [14], but our current results suggest that they might be disease progression factors specifically relevant in the steroid model (Fig. 3C).

It was unfortunate that the samples from two leukopenic mice did not run properly and had to be excluded from the analyses. We acknowledge that this limits the value of some of our results, particularly the comparisons among time-points and the analysis of temporal expression of genes in leukopenic mice. However, as all the infections for the *in vivo* samples of the analysis were carried out simultaneously, we reasoned that repeating just two mice would not provide fully consistent results. Indeed, we tried to include the data obtained from the two leukopenic mice of the preliminary experiment to validate the linear amplification protocol (Fig. 1A), but found that these samples clustered slightly separated from the other *in vivo* samples (Fig S5), and therefore could not be used for the model and temporal analyses. Importantly, including these two mice, the overall profile of expression relative to *in vitro* remained highly similar, such that the 15 genes previously detected to be consistently upregulated *in vivo* VS *in vitro* were still detected (Fig. S5), which shows the strength of the results. In addition, with these two additional mice one more gene was included in the list of candidates, AFUA_5G07210. This gene encodes a homoserine O-acetyltransferase, which has been described as important for virulence in the fungal pathogen *Cryptococcus neoformans* [75].

One of the uncharacterised genes in our list of 15 potentially relevant genes encodes a serine hydroxymethyltransferase (SHMT). SHMT is a pyridoxal phosphate-dependent enzyme of the folate cycle that catalyses the reversible conversion of serine and tetrahydrofolate (THF) into glycine and 5,10-methylenetetrahydrofolate (CH2-THF). Fungi encode two isoforms in the nuclear genome, one cytosolic and one mitochondrial [45]. In *S. cerevisiae* both genes were disrupted singly and in combination, generating auxotrophic viable strains [76]. In contrast to our results, the deletant of the cytosolic SHMT encoding gene (*SHM2*) grew well in the presence of the substrates glycine and formate. In addition, the cytosolic and mitochondrial SHMT encoding genes could also be deleted in the ascomycete *Ashbya gossypii* [77], where supplementation with adenine or glycine could reconstitute growth of the cytosolic *SHMT* (*SHM2*) knock out. Therefore, essentiality of the cytosolic ShmB in *A. fumigatus* suggests that the metabolic implications of disrupting the one carbon-cycle are more complex in this pathogen than in other fungi. Indeed, we previously showed that in contrast to other fungi, the absence of methionine synthase, which forms a junction between the one carbon cycle and the trans-sulfuration pathway, causes an imbalance in cell energetics, possibly initiated by a cellular sensing of purine depletion, that halts fungal growth [40]. Here we found that adenine could partly reconstitute growth of the *shmB_tetOFF* strain in restrictive conditions, which resembles the phenotype we observed with *metH_tetOFF* [40]; but in contrast to the absence of MetH, ATP could not fully reconstitute growth in the absence of ShmB, reflecting that we do not understand the metabolic implications of eliminating the cytosolic serine hydroxymethyltransferase activity. Given its essentiality, we hypothesised that it could constitute a good target for the development of novel antifungals, and indeed we have validated it in a murine model of established infection using the *tetOFF* system. To our knowledge the relevance of serine hydroxymethyltransferases for fungal pathogenicity had not been investigated before. In contrast, this enzyme has been shown to be a promising antimalarial target [78] and implicated in a variety of virulence related traits in bacteria. For instance, a null mutant in *Staphylococcus aureus* showed impaired survival inside macrophages and reduced virulence in a *Galleria mellonella* infection model [79]. In *Helicobacter pylori*, deletion of the SHMT encoding gene *glyA* caused reduced growth rate [80], and in *Pseudomonas aeruginosa* controls rugose colony morphology, increases biofilm formation, abolishes swarming and relates to the redox status of the cells and the control of iron acquisition [81]. Therefore, targeting microbial SMHT enzymes seems to be a promising antimicrobial strategy. Nevertheless, as the current SHMT inhibitors have been developed against the human enzyme, specific inhibitors against the microbial proteins need to be designed. By comparing the pocket landscape of the *A. fumigatus* and human enzymes and performing a docking assay with a known inhibitor of the human SHMT, here we show that there are sufficient differences to pursue the development of a fungal specific agent. A comparable situation occurs in the case of efforts to inhibit the promising antifungal target calcineurin. The human and fungal enzymes are highly similar, such that in this case a drug developed against the human protein (FK506) can inhibit the fungal enzyme and is a potent antifungal agent. Fuelled by structural insight, recent research is focused on developing FK506 fungal specific derivatives, which has already generated promising results [82–84].

In conclusion, in this study we show that *in vivo* transcriptomics is a valid strategy to identify virulence traits in *A. fumigatus*. Using the NanoString technology we have detected 15 genes as potentially relevant for pathogenicity, five of which have already been proven experimentally. We further investigate one novel gene, encoding the cytosolic serine hydroxymethyltransferase, thereby demonstrating that this is an essential gene product for *A. fumigatus* viability and virulence, and that it seems possible to design specific inhibitors for the fungal enzyme. Therefore, we have validated cSHMT as a promising antifungal target for future research efforts.

## MATERIAL AND METHODS

### Fungal strains and culture media

The *Aspergillus fumigatus* strain ATCC46645 [85] was used for the transcriptomic analysis, and the MFIG001 isolate (*ΔKU80*) for genetic manipulation [86]. The strains were routinely grown on Sabouraud solid medium (Oxoid) for 3-5 days at 37 °C for spore harvesting.

For phenotyping analysis, the isolates were inoculated on solid *Aspergillus* Minimal Medium (AMM, 1% glucose, 5 mM ammonium tartrate, 7 mM KCl, 11 mM KH2PO4, 0.25 mM MgSO4, 1× Hutner’s trace elements solution [pH 5.5], 1.5% agar), supplemented as indicated in each section, and incubated at 37 °C for 2-3 days. Phenotypic experiments were performed at least twice independently.

For selection in the presence of resistance markers 50 μg ml^−1^ of hygromycin B or 100 μg ml^−1^ of pyrithiamine (InvivoGen) were added to the AMM in the growth plates.

### Mutant construction

To generate the *ΔshmA* mutant and attempt to construct the *ΔshmB* mutant, we followed a previously optimised protocol [87]. Briefly, upstream and downstream fragments of the genes were amplified by HIFI-PCR and fused to the Hygromycin-B (*hph*) selective cassette by fusion PCR. This deletant cassette was used to replace the target gene by homologous recombination in the *ΔKU80* strain MFIG001.

To construct the *shmB_tetOFF* isolate, we followed CRISPR-Cas9 mediated recombination, as previously described for *A. fumigatus* [88]. Briefly, a repair template containing the *tetOFF* module and a pyrithiamine resistance marker (PtrA), was amplified from pSK606 [39] using Phusion green hot start II high-fidelity PCR master mix (ThermoFisher Scientific). The repair template had 50 bp flanking regions homologous to the upstream and downstream sequences at the ATG start codon, placing the *tetOFF* module immediately 5’- of the *shmB* ORF. The sequence of the targeted gRNA (crRNA) was (GTCCAAATGAAAGAAAACAA|TGG) and was designed using EuPaGDT [89].

To construct the fluorescent SHMT strains, we inserted the correspondent repair template, containing Citrine and the pyrithiamine resistance marker, at the 3’-UTR of each gene using CRISPR-Cas9 as previously descried [88]. Each repair template carried flanking homology regions to the targeted gene. The targeted gRNAs used were (5’-GACAGGCAGGGGGTAGGTGC-3’) and (5’- GGGCACTTTCAGCCTTCCGT-3’) for *shmB* and *shmA*, respectively.

All primers used in the study can be found in Table S3.

### Localisation studies of ShmB-Citrine

*A. fumigatus* conidia were inoculated in 200 μL of filtered autoclaved AMM on 8 well chambers slides (IBIDI) and incubated at 30 °C for 16h. Imagining was performed using a Leica SP8x laser scanning confocal microscope equipped with a 63x (NA 1.4) HC PLAN APO CS2 oil immersion objective. The pinhole was set to one Airy unit and the gain was set to 200. To visualise the localisation of labelled proteins, fluorescence imaging was performed using an excitation of 516 nm and 529 nm emission using a white light laser. Image format captures was 1024x1024 at 400 hz scan speed. Z-stacks of germlings were taken at 1.0 microns and slices were projected as average intensity using ImageJ no adjustments to brightness or contrast were made. Scale bars are shown.

### Murine Infections

Mouse infections experiments were performed under United Kingdom Home Office project license PDF8402B7 and approved by the University of Manchester Ethics Committee and by the Biological Services Facility at the Faculty of Biology, Medicine and Health, University of Manchester.

Outbred CD1 male mice (22 to 26 g) were purchased from Charles Rivers and left to rest for at least one week before the experiment. Mice were allowed access *ad libitum* to water and food throughout the experiment.

For the leukopenic model mice were immunosuppressed with 150 mg/kg of cyclophosphamide (Baxter) on days −3 and −1 plus 1 subcutaneous injection of 250 mg/kg hydrocortisone 21-Acetate (Sigma-Aldrich) on day −1. For the corticosteroid model, mice were injected with a single dose of 40 mg/kg of triamcinolone (Bristol Myers Squibb) on day −1. Bacterial infections were prevented by adding 2 g/L neomycin to the drinking water. Mice were anesthetized by exposure to 2% to 3% isoflurane (Sigma-Aldrich) for 5 to 10 minutes and intranasally infected with 40 μL of a suspension containing 10^5^ spores for the leukopenic model and 10^6^ spores for the corticosteroid model. For both models, 3 mice per time point post infection (16, 24 or 72h) were used. At the designated time-points, mice were sacrificed by a lethal injection of pentobarbital and the lungs were harvested to be immediately snap frozen for RNA isolation.

For the chronic model, we followed a published protocol [25, 90]. Briefly, an immunocompetent mouse was anesthetised by intraperitoneal injection of a ketamine (1%)/xylazine (0,2%) solution and intratracheally inoculated with 50 µL of a suspension of agar beads containing 5×10^7^ conidia/mL. Infection was left to progress for 21 days before the mouse was sacrificed for further processing.

To monitor virulence of the *shmB_tetOFF* strain, we followed our previously optimised protocol [40]. Briefly, groups of 10 mice were infected with 40 μL of a suspension containing 2×10^5^ of the *shmB_tetOFF* or the control *Δcyp51B-cyp51A-tetOFF* strain (inocula were confirmed by plating dilutions of the used suspensions). Five mice per strain received doxycycline (Dox) treatment and five PBS vehicle. Dox treatment started 16h after infection by an intraperitoneal injection (50 mg/kg) and change to Dox-containing food (Envingo Safe-Diet U8200 version 0115 A03 0.625 g/kg Doxycycline Hyclate pellets). Treatment was maintained by subcutaneous 50 mg/kg Dox injections every 12h post infection until the end of the experiment after 72h.

### NanoString analysis: Preparation of samples, hybridisation and data processing

RNA from *in vitro* conditions was isolated as previously described [13]. Briefly, mycelia were shifted from a culture grown overnight in AMM to new flasks containing the defined sources described in Table S2. Mycelia were incubated for 4h in the new conditions, then filtrated trough sterile Miracloth (Merck Millipore) and snap frozen in liquid nitrogen. Mycelia were ground in the constant presence of liquid nitrogen and RNA was isolated using the RNeasy Plant Mini Kit (Qiagen) following the manufacturer’s instructions and including the on column DNAse digestion step. RNA was assessed by a NanoDrop 2000 spectrophotometer (Thermo Fisher Scientific) to determine quality and concentration.

RNA from murine lungs was isolated as follows: Lungs were lyophilized for 48h in a CoolSafe ScanVac freeze drier connected to a VacuuBrand pump and subsequently ground in the presence of liquid nitrogen. The resulting powder was resuspended in TRIzol reagent (Invitrogen – Thermofisher) and processed following the manufacturer’s instructions. RNA was cleaned with Phenol:Chloroform:Isoamyl (25:24:1) (Sigma) and extracted with Chloroform (Invitrogen). Subsequently, RNA was purified using the RNeasy Minikit (Qiagen), following the clean-up protocol, and including the on column DNAse digestion step.

RNA isolated from murine lungs was subjected to the NanoString nCounter Single Cell Gene Expression protocol for linear amplification of target genes. Briefly, RNA was converted to cDNA using SuperScript VILO MasterMix (Life Technologies). To confirm that target RNA was present in the samples, and that cDNA has been properly generated, a Real-Time PCR was performed using the gene AFUA_4G07360 (*metH*) as target, as a RT-PCR for this transcript had been validated before [40]. Genes of interest were linearly amplified using the Multiplex Target Enrichment (MTE) primers (Supplementary File 1) and a PCR protocol consisting of 16 MTE amplification cycles of 94°C 15 seconds and 60°C 4 minutes, with the TaqMan PreAmp Master Mix (Life Technologies).

cDNA samples were incubated for 2 minutes at 94°C and then snap cool on ice prior to addition to the hybridization reaction. Hybridization was performed according to the Elements Tagset Hybridisation setup, as detailed by Nanostring and using the NanoString custom designed probes A and B (Supplementary File 1). The cartridge was read using the flex system, consisting of nCounter Prep Station and the Digital Analyser.

The saw data file (Supplementary File S2) was initially subjected to a Quality Control check. This consisted of the monitorization of the counts of the internal positive control (positive performance for hybridization verification) and the total counts detected for each of the conditions (Fig. S1).

Raw data (Supplementary File 2) was normalised using the nSolver 4.0 software (NanoString). Data was first subjected to Background Subtraction using the geometric mean of the negative control counts. Subsequently, data was normalized using both the geometric mean of positive control count and the geometric mean of the housekeeping genes (Supplementary File 3).

Normalised data was analysed using the Agglomerative Cluster – Heat map option of the nSolver 4.0 software (NanoString). Spearman Correlation was used as distance metric and Average was used as linkage method.

### Computational structure analyses

The crystal structure of the serine hydroxymethyltransferase (SHMT) was obtained from Research Collaboratory for Structural Bioinformatics Protein Data Bank (RCSB PDB, http://www.rcsb.org) in PDB format, referenced by its protein data bank ID 1bj4. The structure of the fungal SHMT was predicted using AlphaFold2 [53]. The amino acid sequence was retrieved from FungiDatabase [91] and utilised as input, with the highest scoring model (pLDDT= 96.9, pTMscore= 0.943) selected for downstream *in silico* analysis.

For the human SHMT, DrugRep [51] and PockDrug [52] were employed for target based binding pocket prediction. DrugRep is a receptor-based screening tool utilising CB-Dock to identify docking pockets of the protein receptor. PockDrug utilizes the fpocket estimation method, based on holo or apo proteins, for predicted pocket estimation. This method involves preliminary detection of cavities capable of binding a ligand of sufficient size, though without ligand proximity information. Pockets with a predicted druggability score above 0.5 from DrugRep were recorded and cross-referenced with suggestions from PockDoc to determine the ’optimal’ binding pocket for subsequent *in-silico* docking.

Ligand Hit-1 was drawn using Marvin JS (https://marvinjs-demo.chemaxon.com/latest/demo.html [92]). The binding mode of Hit-1 to optimal binding pocket in SHMT was predicted using the EADock DS based SwissDock (http://www.swissdock.ch/docking# [55]). The binding modes with the most favourable energies were evaluated with FACTS (Fast Analytical Continuum Treatment of Solvation [93]) and then output as clusters. The best docking pose, exhibiting maximal hydrophobicity pockets coverage and hydrogen bonding, was manually selected. Outputs from all programmes were visualised in PyMOL 2.5 (Schrödinger, LLC).

## ACKOWLEDGEMENTS

We would like to specially thank Prof Sven Krappmann, who guided JA to discover the NanoString technology some years ago and so inspired this work. In addition, we would like to thank Emilia Mellado and Sven Krappmann for critical reading of the manuscript. We acknowledge the use of the Genomic Technologies Facility (Faculty of Biology Medicine and Health, University of Manchester) for the NanoString hybridisation and data read. Help and support from members of the Manchester Fungal Infection Group (MFIG) and the Spanish National Centre for Mycology (LRIM) is greatly appreciated.

**A)** R. Alharthi was funded by the Ministry of Education of Saudi Arabia. J Amich was funded by an MRC Career Development Award (MR/N008707/1). M. Bromley was supported by Wellcome Trust (grants: 219551/Z/19/Z and 208396/Z/17/Z).

## SUPPLEMENTARY FIGURES

**Fig. S1. Quality Control of NanoString run**

**A)** Total number of positive counts (included in the codeset) detected in each condition. **B)** Total number of counts (excluding positive and negative controls) in each condition. All conditions reached the required thresholds except the samples Leukopenic 1.2 and leukopenic 1.3, which were eliminated from all analyses.

**Fig. S2. Temporal dynamics of gene expression**

Fold change of all genes in each *in vitro* condition versus each of the **A)** leukopenic samples and **B)** corticosteroid ordered by the time after infection identified temporal dynamics in the expression of certain genes.

**Fig. S3. Shm supplementary data**

**A)** Alignment of the amino acid sequences of the two serine hydroxymethyltransferase enzymes encoded in *A. fumigatus* genome. The proteins have 61.9% identity and 79.9% similarity. ShmA has a long N-terminus region that is absent in ShmB and which after further analysis (see text) suggests that ShmA is mitochondrial. **B)** Alignment of the amino acid sequences of *A. fumigatus* (ShmA and ShmB) and *S. cerevisiae* (SHM1 and SHM2) serine hydroxymethyltransferase proteins. As the yeast mitochondrial enzyme SHM1, ShmA shows an elongated N-terminus, suggesting that this is the mitochondrial enzyme in *A. fumigatus*. **C)** Two independent transformants of *ΔshmA* grew normally on both Sabouraud (SAB) rich and minimal (MM) media, proving that this gene is not essential. **D)** Transcription of *shmB* in the *shmB_tetOFF* strain decreased very strongly upon Dox addition. Graph depicts the mean and the SD. Data was analysed with one-way ANOVA with Dunnett’s multiple comparisons, *p* <0.001.

**Fig. S4. Colistin permeabilises the membrane of *A. fumigatus* swollen conidia**

Swollen conidia are impermeable to the dyes FITC and Zombie-Aqua^TM^. However, when germinated in the presence of 4 µg/mL colistin these dyes were able to penetrate inside the cells, demonstrating that the membrane has been permeabilised. A) Image of the central plane of a z-stack covering the full field of view. B) Zoom of individual conidia at their own central plane.

**Fig. S5. Hierarchical clustering with two additional mice**

The inclusion of two extra mice (from the experiment described in Figure 1A) did not substantially modified the expression profile, although the two mice clustered separated from the rest (purple arrow). There was still a complete separation of *in vitro* and *in vivo* samples, and the same 15 genes identified before as always expressed at higher levels (red line) were maintained. Additionally, one more gene got included in this list, AFUA_5G07210 (red arrow), encoding a homoserine O- acetyltransferase.

## SUPPLEMENTARY FILES

**Table S1**: Genes included in the NanoString custom codeset.

**Table S2**: *In vitro* and *in vivo* conditions assayed for the NanoString run.

**Table S3**: Primers employed in this study.

**Supplementary File 1**: NanoString designed Probes A and B for hybridisation and MTE primers for enrichment.

**Supplementary File 2**: Raw data from digital analysis of hybridised codesets.

**Supplementary File 3:** Normalized data used for the analyses.

