## Supplementary Figures for "The sulfur-related metabolic status of *Aspergillus fumigatus* during infection reveals cytosolic serine hydroxymethyltransferase as a promising antifungal target"

Fig. S1

A

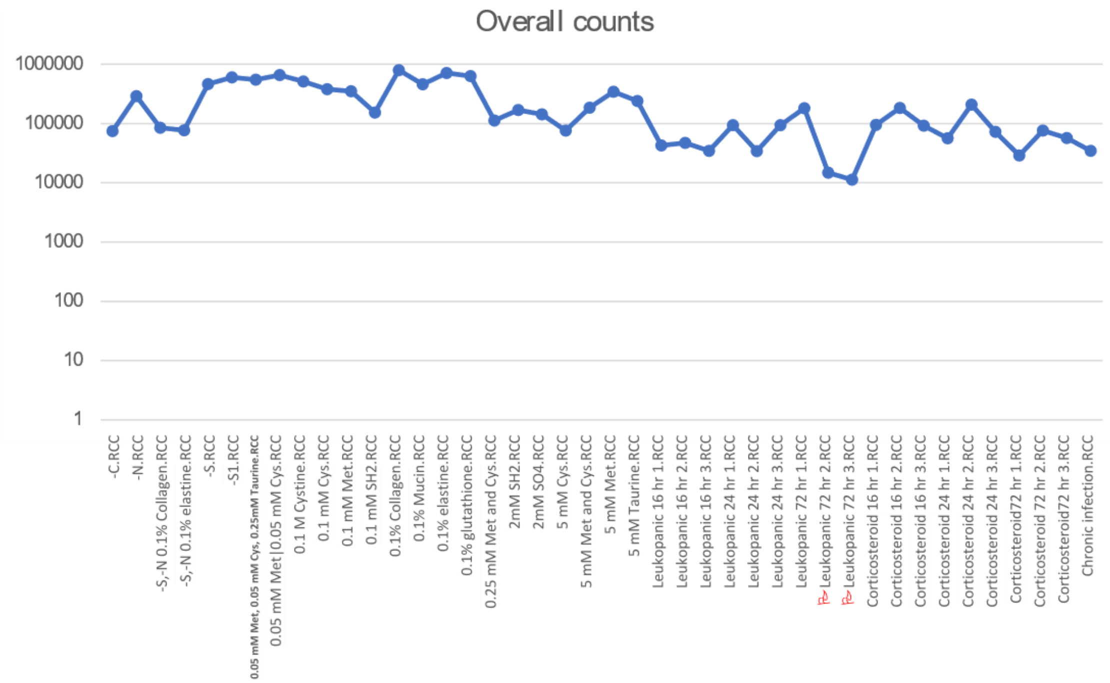
Overall counts
1000000
100000
10000
1000
100
10
1
-C.RCC
-N.RCC
-S,-N 0.1% Collagen.RCC
-S,-N 0.1% elastine.RCC
-S.RCC
-S1.RCC
0.05 mM Met, 0.05 mM Cys, 0.25mM Taurine.RCC
0.05 mM Met|0.05 mM Cys.RCC
0.1 M Cystine.RCC
0.1 mM Cys.RCC
0.1 mM Met.RCC
0.1 mM SH2.RCC
0.1% Collagen.RCC
0.1% Mucin.RCC
0.1% elastine.RCC
0.1% glutathione.RCC
0.25 mM Met and Cys.RCC
2mM SH2.RCC
2mM SO4.RCC
5 mM Cys.RCC
5 mM Met and Cys.RCC
5 mM Met.RCC
5 mM Taurine.RCC
Leukopanic 16 hr 1.RCC
Leukopanic 16 hr 2.RCC
Leukopanic 16 hr 3.RCC
Leukopanic 24 hr 1.RCC
Leukopanic 24 hr 2.RCC
Leukopanic 24 hr 3.RCC
Leukopanic 72 hr 1.RCC
Leukopanic 72 hr 2.RCC
Leukopanic 72 hr 3.RCC
Corticosteroid 16 hr 1.RCC
Corticosteroid 16 hr 2.RCC
Corticosteroid 16 hr 3.RCC
Corticosteroid 24 hr 1.RCC
Corticosteroid 24 hr 2.RCC
Corticosteroid 24 hr 3.RCC
Corticosteroid72 hr 1.RCC
Corticosteroid 72 hr 2.RCC
Corticosteroid72 hr 3.RCC
Chronic infection.RCC

B

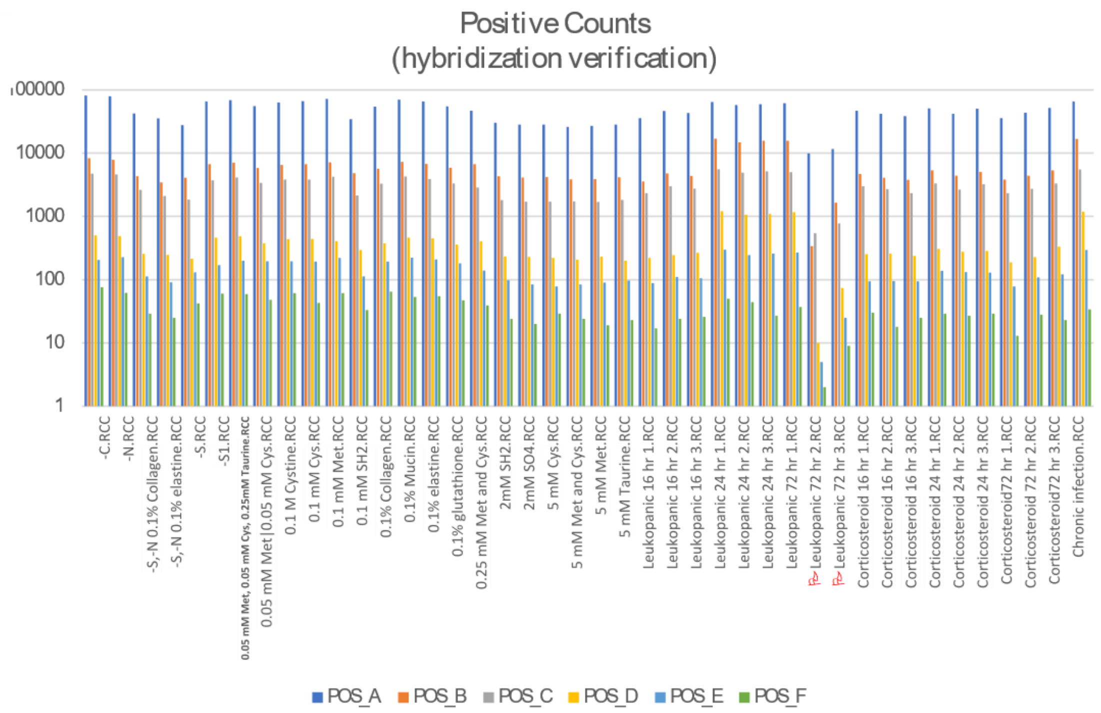
Positive Counts
(hybridization verification)
100000
10000
1000
100
10
1
-C.RCC
-N.RCC
-S,-N 0.1% Collagen.RCC
-S,-N 0.1% elastine.RCC
-S.RCC
-S1.RCC
0.05 mM Met, 0.05 mM Cys, 0.25mM Taurine.RCC
0.05 mM Met|0.05 mM Cys.RCC
0.1 M Cystine.RCC
0.1 mM Cys.RCC
0.1 mM Met.RCC
0.1 mM SH2.RCC
0.1% Collagen.RCC
0.1% Mucin.RCC
0.1% elastine.RCC
0.1% glutathione.RCC
0.25 mM Met and Cys.RCC
2mM SH2.RCC
2mM SO4.RCC
5 mM Cys.RCC
5 mM Met and Cys.RCC
5 mM Met.RCC
5 mM Taurine.RCC
Leukopanic 16 hr 1.RCC
Leukopanic 16 hr 2.RCC
Leukopanic 16 hr 3.RCC
Leukopanic 24 hr 1.RCC
Leukopanic 24 hr 2.RCC
Leukopanic 24 hr 3.RCC
Leukopanic 72 hr 1.RCC
Leukopanic 72 hr 2.RCC
Leukopanic 72 hr 3.RCC
Corticosteroid 16 hr 1.RCC
Corticosteroid 16 hr 2.RCC
Corticosteroid 16 hr 3.RCC
Corticosteroid 24 hr 1.RCC
Corticosteroid 24 hr 2.RCC
Corticosteroid 24 hr 3.RCC
Corticosteroid72 hr 1.RCC
Corticosteroid 72 hr 2.RCC
Corticosteroid72 hr 3.RCC
Chronic infection.RCC
POS_A
POS_B
POS_C
POS_D
POS_E
POS_F

Fig. S2

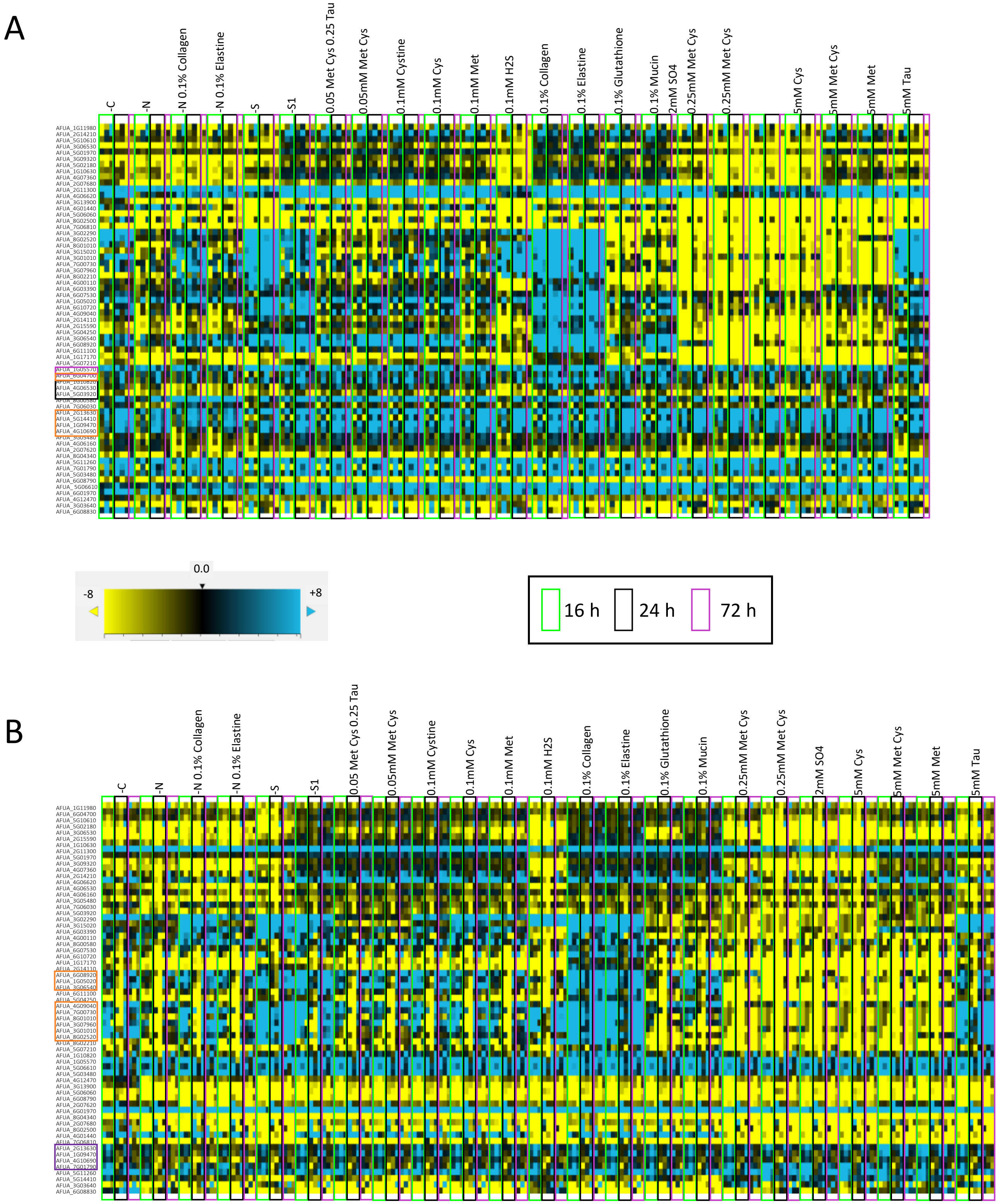
A
-C
-N
-N 0.1% Collagen
-N 0.1% Elastine
-S
-S1
0.05 Met Cys 0.25 Tau
0.05mM Met Cys
0.1mM Cystine
0.1mM Cys
0.1mM Met
0.1mM H2S
0.1% Collagen
0.1% Elastine
0.1% Glutathione
0.1% Mucin
2mM SO4
0.25mM Met Cys
0.25mM Met Cys
5mM Cys
5mM Met Cys
5mM Met
5mM Tau
0.0
-8
+8
16 h
24 h
72 h
B
-C
-N
-N 0.1% Collagen
-N 0.1% Elastine
-S
-S1
0.05 Met Cys 0.25 Tau
0.05mM Met Cys
0.1mM Cystine
0.1mM Cys
0.1mM Met
0.1mM H2S
0.1% Collagen
0.1% Elastine
0.1% Glutathione
0.1% Mucin
0.25mM Met Cys
0.25mM Met Cys
2mM SO4
5mM Cys
5mM Met Cys
5mM Met
5mM Tau

### Fig. S3

A

```
shmB ---------------------------------------------------MATYALSQAH  10
shmA MSLSRCGRQALRLIPRSGSSSRATTVITTARHPGFQLQTPTAASRNVQWRSVSSSSRDGQ  60
                                                     .: :  :.:

shmB REQMEKSLVDSDPEIAQIMEKEIQRQRESILLIASENVTSRAVFDALGSPMSNKYSEGYP  70
shmA QHLLSASLEEQDPTVYNILQKEKKRQKHFINLIPSENFTSQAVLDALGSVMQNKYSEGYP  120
     :. :. ** :.** : :*::** :**:. * ** ***.**:**:***** *.********

shmB GARYYGGNQHIDAIELTCQARALKAFNLDPEKWGVNVQCLSGSPANLEVYQALMRPHDRL  130
shmA GARYYGGNEFIDESERLCQQRALETFRLDPEEWGVNVQALSGSPANLYAISAVLNTHDRL  180
     ********:.**  *  ** ***::*.****:******.********  . .*::. ****

shmB MGLDLPHGGHLSHGYQTPSRKISAVSTYFETFPYRVNTETGIIDYDTLEANAELYRPKCL  190
shmA MGLDLPHGGHLSHGYQTPTKKISFISKYFETLPYRLDESTGLIDYDGAEKLALLYRPKLI  240
     ******************::*** :*.****:***:: .**:****  *  * ***** :

shmB VAGTSAYCRLIDYGRMRKIADKVGAYLIVDMAHISGLVAAGVIPSPFEYADVVTTTTHKS  250
shmA IAGTSAYSRLIDYPRMRQIADAAGAYLLSDMAHISGLVAAGVLPSPFPHSDIVTTTTHKS  300
     :******.***** ***:*** .****: *************:****  ::*:********

shmB LRGPRGAMIFFRKGVRSTDPKTGKEIMYDLEGPINFSVFPGHQGGPHNHTITALAVALKQ  310
shmA LRGPRGAMIFYRKGVRRTDKK-GNKEMYDLENLINASVFPGHQGGPHNHTITALSVALKQ  359
     **********:****** ** * *:: *****. ** ******************:*****

shmB AATPEFRQYQEQVLKNAKALEVE------FKALGHKLVSDGTDSHMVLLDLRPKGLDGAR  364
shmA AQTPEFKAYQETVLANAKALSERLGGPINNGGLGYNIVSGGTDNHLVLVDLKNRGVDGAR  419
     * ****: *** ** *****. .         .**:::**.***.*:**:**: :*:****

shmB VEAVLEQINIACNKNSIPGDKSALTPCGIRIGTPAMTSRGMSEEDFKRVARYIDQVINLC  424
shmA VERVLELCGVASNKNTVPGDQSALKPGGLRLGTPAMTTRGFQPEDFRRVADIVDRAVIIT  479
     ** ***  .:*.***::***:***.* *:*:******:**:. ***:***  :*:.: :

shmB KSIQADL--------PKEANKLKDFKAKVA-SGSVPEILALRKEVAEWASTYPLPV--   471
shmA QKLDKAAKESAAAKGVKNPNTVKAFLDYVGEGDEISEIVLLRKEVEDWVGTFSLPWDD   537
     :.::            *: *.:* *   *. ...: **: ***** :*..*: **
```

B

```
SHM1   ------------MFPRASAL----------------------AKCMATVHRRGLLTSGA  25
shmA   MSLSRCGRQALRLIPRSGSSSRATTVITTARHPGFQLQTPTAASRNVQWRSVSSSSRDGQ  60
shmB   ---------------------------------------------------MATYALSQAH  10
SHM2   ----------------------------------------------------MPYTLSDAH  9
                                                            ..

SHM1   QSLVSKPVSEGDPEMFDILQQERHRQKHSITLIPSENFTSKAVMDLLGSELQNKYSEGYP  85
shmA   QHLLSASLEEQDPTVYNILQKEKKRQKHFINLIPSENFTSQAVLDALGSVMQNKYSEGYP  120
shmB   REQMEKSLVDSDPEIAQIMEKEIQRQRESILLIASENVTSRAVFDALGSPMSNKYSEGYP  70
SHM2   HKLITSHLVDTDPEVDSIIKDEIERQKHSIDLIASENFTSTSVFDALGTPLSNKYSEGYP  69
       :  :   : : ** : .*::.* .**:. * ** ***.** :*:* **: :.********

SHM1   GERYYGGNEIIDKSESLCQARALELYGLDPAKWGVNVQPLSGAPANLYVYSAIMNVGERL  145
shmA   GARYYGGNEFIDESERLCQQRALETFRLDPEEWGVNVQALSGSPANLYAISAVLNTHDRL  180
shmB   GARYYGGNQHIDAIELTCQARALKAFNLDPEKWGVNVQCLSGSPANLEVYQALMRPHDRL  130
SHM2   GARYYGGNEHIDRMEILCQQRALKAFHVTPDKWGVNVQTLSGSPANLQVYQAIMKPHERL  129
       * ******: **  *  ** ***: : : * :****** ***:**** . .*::.  :**

SHM1   MGLDLPDGGHLSHGYQLKSGTPISFISKYFQSMPYHVDHTTGLIDYDNLQVLAKAFRPKV  205
shmA   MGLDLPHGGHLSHGYQTPT-KKISFISKYFETLPYRLDESTGLIDYDGAEKLALLYRPKL  239
shmB   MGLDLPHGGHLSHGYQTPS-RKISAVSTYFETFPYRVNTETGIIDYDTLEANAELYRPKC  189
SHM2   MGLYLPDGGHLSHGYATEN-RKISAVSTYFESFPYRVNPETGIIDYDTLEKNAILYRPKV  188
       *** **.********    .   ** :*.**:::**::: **:****  :  *  :***

SHM1   IVAGTSAYSRLIDYARFKEISQGCGAYLMSDMAHISGLVAANVVPSPFEHSDIVTTTTHK  265
shmA   IIAGTSAYSRLIDYPRMRQIADAAGAYLLSDMAHISGLVAAGVLPSPFPHSDIVTTTTHK  299
shmB   LVAGTSAYCRLIDYGRMRKIADKVGAYLIVDMAHISGLVAAGVIPSPFEYADVVTTTTHK  249
SHM2   LVAGTSAYCRLIDYKRMREIADKCGAYLMVDMAHISGLIAAGVIPSPFEYADIVTTTTHK  248
       ::******.***** *:::*::  ****: ********:**.*:**** ::*:*******

SHM1   SLRGPRGAMIFFRKGIKSVTKK-GKEIPYELEKKINFSVFPGHQGGPHNHTIGAMAVALK  324
shmA   SLRGPRGAMIFYRKGVRRTDKK-GNKEMYDLENLINASVFPGHQGGPHNHTITALSVALK  358
shmB   SLRGPRGAMIFFRKGVRSTDPKTGKEIMYDLEGPINFSVFPGHQGGPHNHTITALAVALK  309
SHM2   SLRGPRGAMIFFRRGVRSINPKTGKEVLYDLENPINFSVFPGHQGGPHNHTIAALATALK  308
       ***********:*:*::    * *::  *:**  ** *************** *::.***

SHM1   QAMSPEFKEYQQKIVDNSKWFAQEL------TKMGYKLVSGGTDNHLIVIDLSGTQVDGA  378
shmA   QAQTPEFKAYQETVLANAKALSERLGGPINNGGLGYNIVSGGTDNHLVLVDLKNRGVDGA  418
shmB   QAATPEFRQYQEQVLKNAKALEVEF------KALGHKLVSDGTDSHMVLLDLRPKGLDGA  363
SHM2   QAATPEFKEYQTQVLKNAKALESEF------KNLGYRLVSNGTDSHMVLVSLREKGVDGA  362
       ** :***: ** :: *:* : .:       :*:.:**.***.*:::::.*    :***

SHM1   RVETILSALNIAANKNTIPGDKSALFPSGLRIGTPAMTTRGFGREEFSQVAKYIDSAVKL  438
shmA   RVERVLELCGVASNKNTVPGDQSALKPGGLRLGTPAMTTRGFQPEDFRRVADIVDRAVII  478
shmB   RVEAVLEQINIACNKNSIPGDKSALTPCGIRIGTPAMTSRGMSEEDFKRVARYIDQVINL  423
SHM2   RVEYICEKINIALNKNSIPGDKSALVPGGVRIGAPAMTTRGMGEEDFHRIVQYINKAVEF  422
       *** : .  .:* ***::***:*** * *:*:*:****:**:  *:* ::.  :: .: :

SHM1   AENLKTLEPTT----KLDARSRLNEFKKLCNESS---EVAALSGEISKWVGQYPVPGDI   490
shmA   TQKLDKAAKESAAAKGVKNPNTVKAFLDYVGEGDEISEIVLLRKEVEDWVGTFSLPWDD   537
shmB   CKSIQADLP--------KEANKLKDFKAKVASGS-VPEILALRKEVAEWASTYPLPV--   471
SHM2   AQQVQQSLP--------KDACRLKDFKAKVDEGS-D-VLNTWKKEIYDWAGEYPLAV--   469
        :.:.           .    :: *     ...    :     *: .*.. : :
```

C

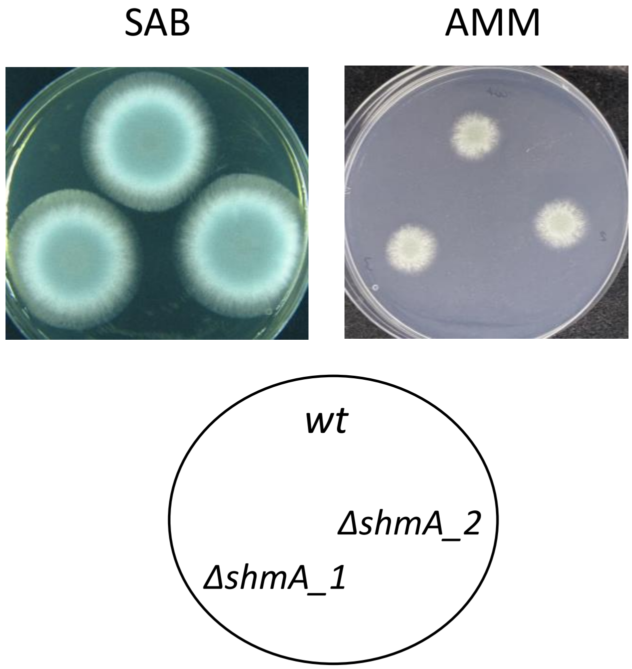


D

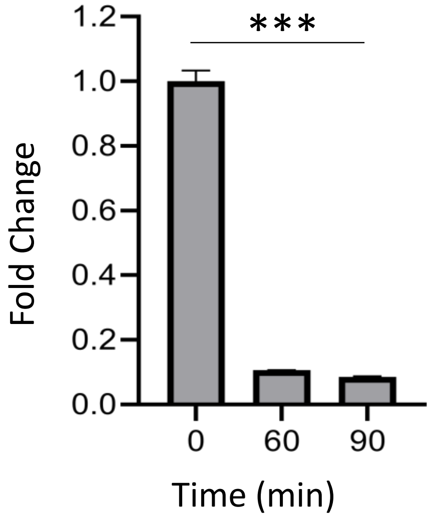

Fig. S4

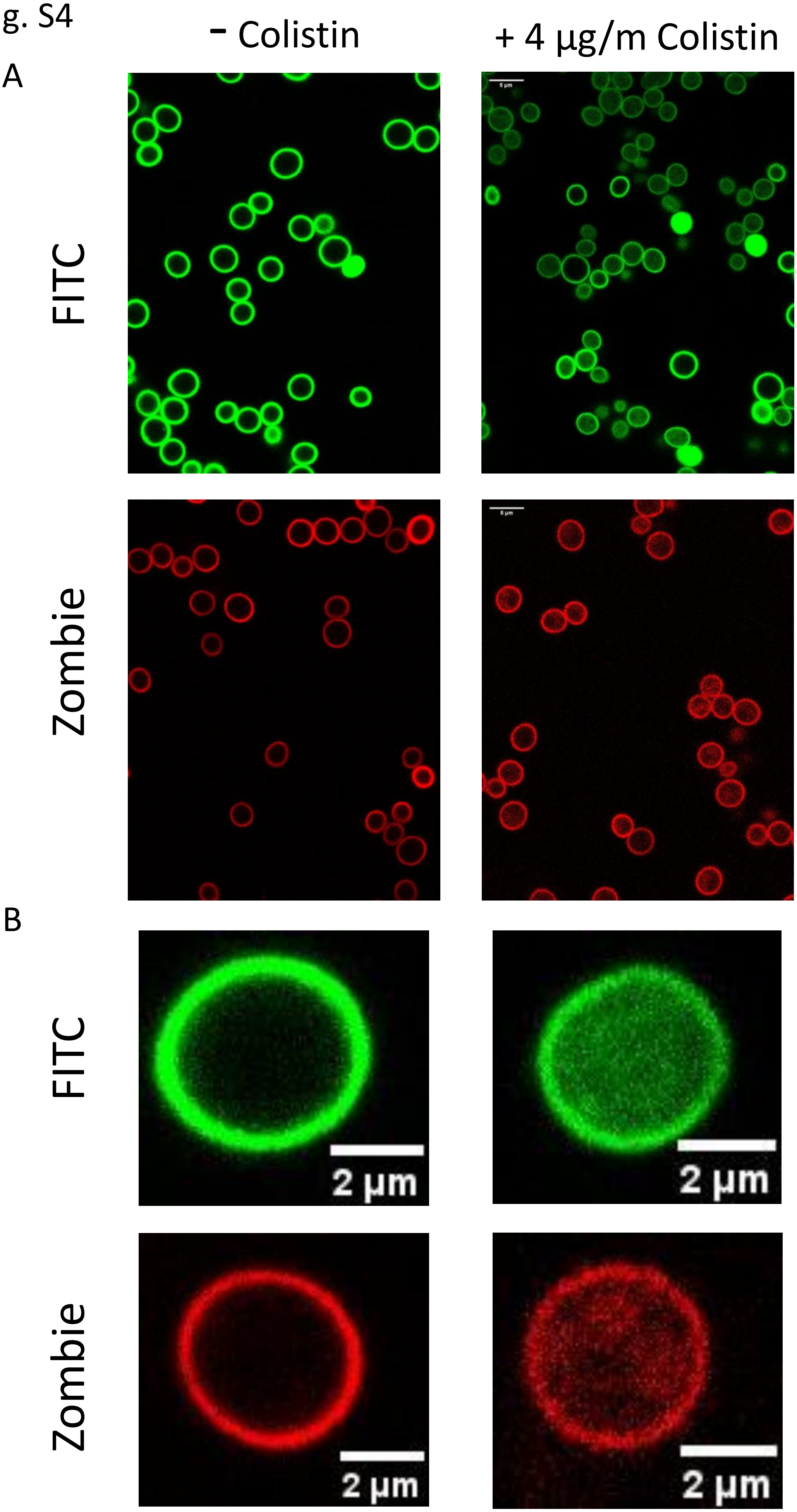
- Colistin
+ 4 µg/m Colistin
A
FITC
Zombie
B
FITC
Zombie
2 µm
2 µm
2 µm
2 µm

Fig. S5

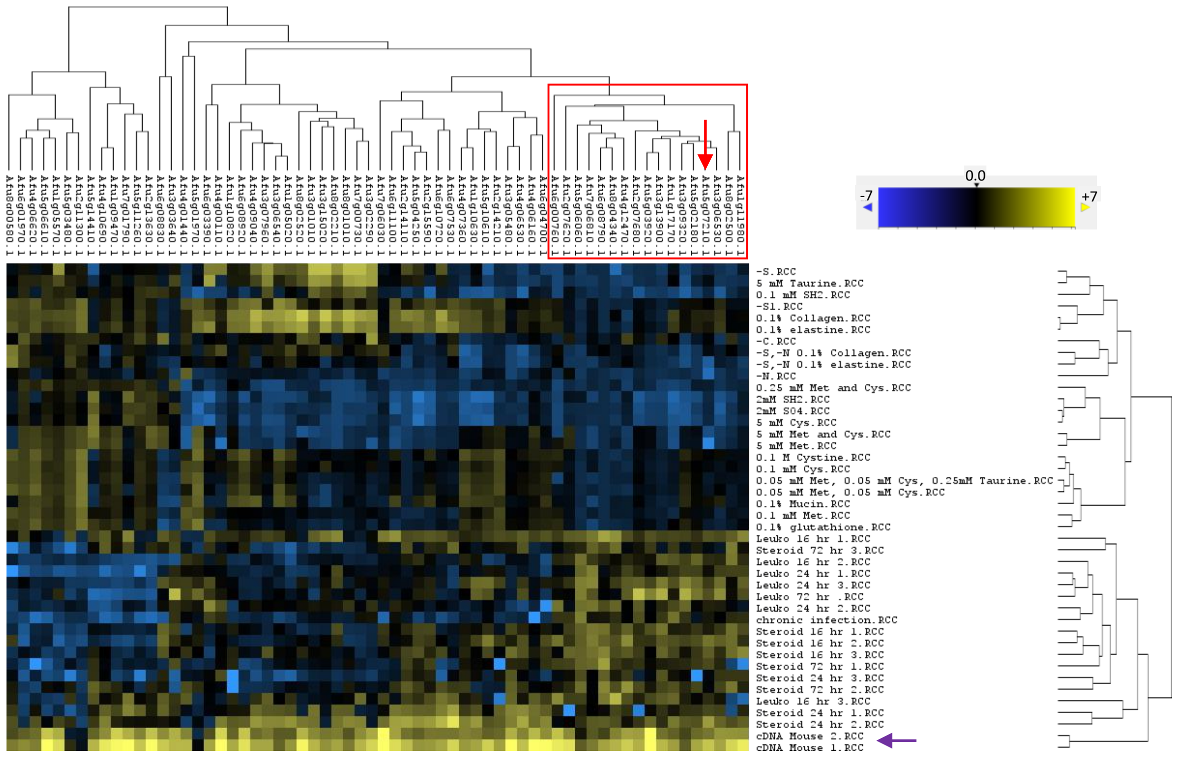

Afu8g00580.1
Afu6g01970.1
Afu4g06620.1
Afu5g06610.1
Afu1g05570.1
Afu5g03480.1
Afu2g11300.1
Afu5g14410.1
Afu4g10690.1
Afu1g09470.1
Afu7g01790.1
Afu5g11260.1
Afu2g13630.1
Afu6g08830.1
Afu3g03640.1
Afu4g01440.1
Afu5g01970.1
Afu6g03390.1
Afu4g00110.1
Afu1g10820.1
Afu6g08920.1
Afu4g09040.1
Afu3g07960.1
Afu3g06540.1
Afu1g05020.1
Afu8g02520.1
Afu3g01010.1
Afu3g15020.1
Afu8g02210.1
Afu8g01010.1
Afu7g00730.1
Afu3g02290.1
Afu7g06030.1
Afu6g11100.1
Afu2g14110.1
Afu5g04250.1
Afu2g15590.1
Afu6g10720.1
Afu6g07530.1
Afu4g07360.1
Afu1g10630.1
Afu5g10610.1
Afu2g14210.1
Afu3g05480.1
Afu4g06530.1
Afu4g06160.1
Afu6g04700.1
Afu6g00760.1
Afu2g07620.1
Afu5g06060.1
Afu7g06810.1
Afu6g08790.1
Afu8g04340.1
Afu4g12470.1
Afu2g07680.1
Afu5g03920.1
Afu3g13900.1
Afu1g17170.1
Afu3g09320.1
Afu5g02180.1
Afu5g07210.1
Afu3g06530.1
Afu8g02500.1
Afu1g11980.1
0.0
-7
+7
-S.RCC
5 mM Taurine.RCC
0.1 mM SH2.RCC
-S1.RCC
0.1% Collagen.RCC
0.1% elastine.RCC
-C.RCC
-S,-N 0.1% Collagen.RCC
-S,-N 0.1% elastine.RCC
-N.RCC
0.25 mM Met and Cys.RCC
2mM SH2.RCC
2mM SO4.RCC
5 mM Cys.RCC
5 mM Met and Cys.RCC
5 mM Met.RCC
0.1 M Cystine.RCC
0.1 mM Cys.RCC
0.05 mM Met, 0.05 mM Cys, 0.25mM Taurine.RCC
0.05 mM Met, 0.05 mM Cys.RCC
0.1% Mucin.RCC
0.1 mM Met.RCC
0.1% glutathione.RCC
Leuko 16 hr 1.RCC
Steroid 72 hr 3.RCC
Leuko 16 hr 2.RCC
Leuko 24 hr 1.RCC
Leuko 24 hr 3.RCC
Leuko 72 hr .RCC
Leuko 24 hr 2.RCC
chronic infection.RCC
Steroid 16 hr 1.RCC
Steroid 16 hr 2.RCC
Steroid 16 hr 3.RCC
Steroid 72 hr 1.RCC
Steroid 24 hr 3.RCC
Steroid 72 hr 2.RCC
Leuko 16 hr 3.RCC
Steroid 24 hr 1.RCC
Steroid 24 hr 2.RCC
cDNA Mouse 2.RCC
cDNA Mouse 1.RCC
