## Supplementary material for "The sulfur-related metabolic status of *Aspergillus fumigatus* during infection reveals cytosolic serine hydroxymethyltransferase as a promising antifungal target": Table S3

**Table S3**. Primers used in this study

|  | Description | Sequence 5’ 🡪 3’ |
| --- | --- | --- |
| shmA deletion AFUA_2G07810 | P1 | GTTCGCATGCCTGTTTGAGG |
|  | P2 | TAGTTCTGTTACCGAGCCGGTGTCAGGAAAAACGGGGTGA |
|  | P3 | GCTCTGAACGATATGCTCCAACGCTTCTTTTGGCGGGTTAGC |
|  | P4 | AGCTCGTCACTTGTGAGCTG |
|  | P5 | TCACGTTGCAGAGTCCTGAC |
|  | P6 | GCGTTATACGGATCGCCCA |
| smhB deletion (attempt) AFUA_3G09320 | P1 | TGGCCCTAAGAGTATCGCCT |
|  | P2 | TAGTTCTGTTACCGAGCCGGGAGAAGGGGAGAGAGGAGGA |
|  | P3 | GCTCTGAACGATATGCTCCAACCAGGCTTTTGACGAGTTCAACA |
|  | P4 | CCCCTACGTCAAGCTCCATG |
|  | P5 | CCCAGACTCTCGTTGGTGTC |
|  | P6 | GTTCTCACGACTCTTCCCCG |
| gRNA (tetOFF) | Tet-OFF Fw | GCCCAACCTTTTTTCCTCCTCTCTCCCCTTCTCTCTTCATCCCTATCATCTTCTAGAATGCCCCACCGTT |
|  | Tet-OFF Rv | GTGGGATGATTCACCTCACGGTGAGCCTGCGACAAAGCGTAGGTGGCCATGGTGATGTCTGCTCAAGCG |
| tetOFF validation | Fw 1 | TTCGACGGAAGACTATCTCG |
|  | Rv 1 | ACGTTCTCAGAGGCAATCAG |
|  | Fw 2 | AGGCCCTTTCGTCTTCACTC |
|  | Rv 2 | ACCACCGTAGTAACGAGCAC |
| shmB RT-PCR | Fw | CGTCTCATGGGTCTGGATCT |
|  | Rv | TGCCGGTCTCAGTGTTAACA |
| Mcitrine insertion | shmB gRNA | GACAGGCAGGGGGTAGGTGC |
|  | shmB tetOFF Fw | CTCTCCGCAAGGAGGTCGCCGAGTGGGCTAGCACCTACCCCCTGCCTGTCATGGTCTCCAAGGGCGAGGA |
|  | shmB tetOFF Rv | AGAAAAGCATATTTTGGTATCAAGTGAGATTATTTGTTCTGCCCTGATGCGCCAATTGATTACGGGATCC |
|  | shmA gRNA | GGGCACTTTCAGCCTTCCGT |
|  | shmA tetOFF Fw | GAAAGGAGGTGGAAGACTGGGTGGGCACTTTCAGCCTTCCGTGAGATGATATGGTCTCCAAGGGCGAGGA |
|  | shmA tetOFF Rv | GAAGCCCTCAAAACATCATCTTTTCATTAAATAGCGGCAATGTATACTTTGCCAATTGATTACGGGATCC |
| Mcitrine insertion validation | shmB tetOFF Fw | GTATGAGCGAGGAGGACTTC |
|  | shmB tetOFF Rv | TGAAGATGAGAAGCTGGGAG |
|  | shmA tetOFF Fw | TCCTTGAACTCTGTGGTGTC |
|  | shmA tetOFF Rv | CAAATAAACAGCCTCACCCG |
